# Beyond the Brake: the Subthalamic Nucleus Predominantly Facilitates Action in Non-human Primates

**DOI:** 10.64898/2025.12.09.692702

**Authors:** Atsushi Yoshida, Richard J. Krauzlis, Okihide Hikosaka

## Abstract

The subthalamic nucleus (STN) is a core component of the basal ganglia circuitry, traditionally viewed as a “brake” that suppresses motor output via the indirect and hyperdirect pathways. While this model explains certain aspects of motor control, it fails to account for the complex involvement of the STN in dynamic, value-based behaviors. Movement-related increases in STN neuronal activity have been well documented across species, but the functional organization of these responses during value-based decisions has remained unclear. Here, we recorded 187 STN neurons from macaque monkeys performing a value-based sequential choice task. Data-driven clustering identified three functionally distinct populations. Two populations (comprising approximately 89% of task-responsive neurons) exhibited facilitative activity modulated by learned object value. Temporal alignment analysis revealed that the majority of these neurons were target-locked, suggesting a predominantly stimulus-driven evaluation signal rather than a strictly movement-locked command. A third, smaller population encoded a negative value signal upon presentation of unrewarded objects, independent of subsequent motor strategy. Response latencies of the facilitative STN populations systematically preceded those of functionally analogous populations in the GPe recorded during the same task paradigm, supporting an STN-GPe facilitative pathway for value-based action selection. These findings reveal a functional organization within the ventral STN in which evaluation of stimulus target value, beyond the traditional motor suppressive model, is the predominant mode of processing.

## Introduction

The subthalamic nucleus (STN) is a crucial component of the basal ganglia circuitry that has traditionally been viewed as responsible for suppressing motor output, a concept rooted in classical models of the indirect and hyperdirect pathways.^1–10^ This “brake” model is supported by extensive anatomical, physiological, and lesion studies. However, this classical view is increasingly challenged by emerging evidence suggesting a more complex and multifaceted role.

Indeed, movement-related increases in STN neuronal activity have been a consistent finding across recording studies. Early studies in non-human primates reported excitatory responses in STN neurons associated with limb movements,^11,12^ and subsequent work showed that the majority of STN neurons modulate their activity in relation to active movement in both healthy and parkinsonian states.^13^ More recent studies have reinforced the predominance of facilitatory-type responses across species and task contexts. For example, in rodents performing a stop-signal task, STN neurons related to movement cancellation were identified, but motor-related neurons were more prevalent.^14^ Similarly, STN neurons in rodents increased their activity at the onset of and during locomotion, and optogenetic inhibition of this activity disrupted locomotor rhythms.^15^ In non-human primates performing go/no-go and stop-signal tasks, STN activity was associated with both motor execution and cancellation.^16,17^ Beyond these single-neuron recording studies, optogenetic studies in rodents demonstrate that STN stimulation can elicit rapid movements,^15,18^ human neuroimaging reveals that STN activity dynamically adapts to behavioral context,^19,20^ and clinical findings from deep brain stimulation in Parkinson’s disease show complex effects on both motor control and impulsivity.^21–23^ Taken together, these findings suggest that the role of the STN extends far beyond simple motor suppression.

What remains unclear, however, is how these facilitative responses are functionally organized during value-based decision-making, where the STN must integrate stimulus evaluation with action selection. Prior studies have largely employed reactive motor tasks (e.g., reaching, grip, stop-signal) that do not manipulate learned stimulus value. Furthermore, whether individual STN neurons encode sensory-driven evaluation signals, motor execution signals, or both has not been systematically examined. Matsumura et al. (1992) distinguished visual and saccade-related responses in the STN during visually guided saccades^24^, but their task did not include a value manipulation, leaving the relationship between temporal alignment and value coding unexplored.

Addressing this question requires consideration of the internal organization of the STN, because the nucleus is not a homogeneous structure. Anatomically, the STN is subdivided into functionally distinct territories that include sensorimotor, oculomotor, associative, and limbic regions, each defined by their cortical and subcortical inputs.^25,26^ These subregions differ in their connectivity and may serve distinct functional roles, raising the possibility that the balance between facilitative and suppressive activity varies across territories. The present study targeted the ventral STN, which overlaps with the oculomotor and associative territories, using a saccade-based task that requires value-based decisions.

A parallel debate surrounds the principal target of the STN, the globus pallidus external segment (GPe). Also traditionally viewed as an inhibitory station, a growing body of evidence now implicates the GPe in action facilitation.^27–30^ In a previous study using the same behavioral paradigm as the present work, we identified a distinct GPe neuronal population that facilitates action initiation and demonstrated that this function causally depends on excitatory inputs.^31^ Given the dense excitatory projection from the STN to the GPe,^25,32^ these findings led us to hypothesize that the STN is a major source of excitatory input to the GPe and may contribute to the excitatory drive underlying this facilitative function.

Here, we recorded from 187 STN neurons while two macaque monkeys performed a value-based sequential choice task. Using data-driven clustering, we identified three functionally distinct populations and characterized their temporal alignment properties, spatial selectivity, and value coding. We compared the response latencies of these populations with functionally analogous populations in the GPe recorded during the identical behavioral paradigm.^31^ Our results reveal a functional organization within the ventral STN in which the predominant mode of processing is stimulus-driven value evaluation that facilitates action selection, rather than direct motor facilitation or suppression.

## Results

### Behavioral performance

The two monkeys (C and S) performed a sequential choice task (Figure 1A) in which they decided whether to accept a “good” (rewarding) object or reject a “bad” (non-rewarding) object. To reject a bad object, they could employ one of three strategies. They could generate a “return” saccade back to fixation after briefly glancing at the bad object, “stay” at the central fixation point without looking, or look at “other” locations in the visual scene (Figure 1B). During each recording session, one of six different sets of objects and scenes was randomly selected (Figure 1C). To dissociate neural activity related to learned value from responses to simple visual features, the design included objects with stable values as well as a “flexible value” condition. In this flexible condition (Scenes 3 and 4), the reward contingencies for the same pair of objects were reversed, requiring the monkeys to adapt their choices based on the scene context.

**Figure 1.**
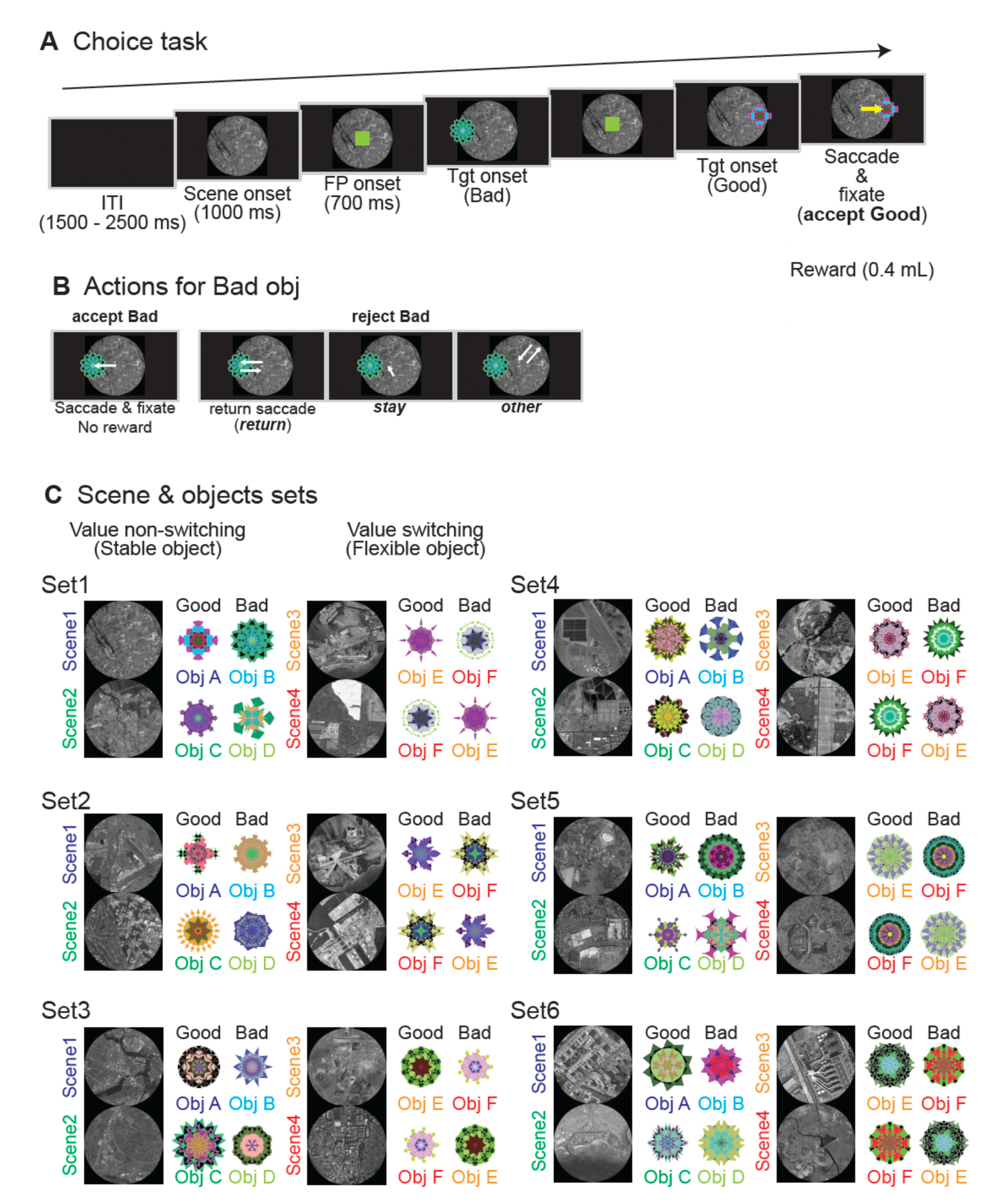
The sequential choice task. (A) Trial sequence. Each trial began with the presentation of a background scene (1000 ms) and a central fixation point (FP, 700 ms). A single object, either “good” (rewarded) or “bad” (non-rewarded), was then presented at one of six peripheral locations. Monkeys received a liquid reward for making a saccade to and fixating on a good object. (B) Response options for bad objects. When a bad object was presented, monkeys could reject it using one of three strategies: a “return” saccade (looking at the object only briefly before making a saccade back to the center), “stay” (maintaining central fixation), or “other” (looking away from both the object and center). Incorrectly accepting a bad object resulted in no reward. (C) Context-dependent value assignment. Six different sets of objects and scenes were used across sessions. Within each set, Scenes 1 and 2 featured objects with stable values. In Scenes 3 and 4, the same pair of objects was used, but their reward contingencies were reversed (“flexible value”), requiring the monkeys to adapt their choices based on the scene context.

**Figure S1.**
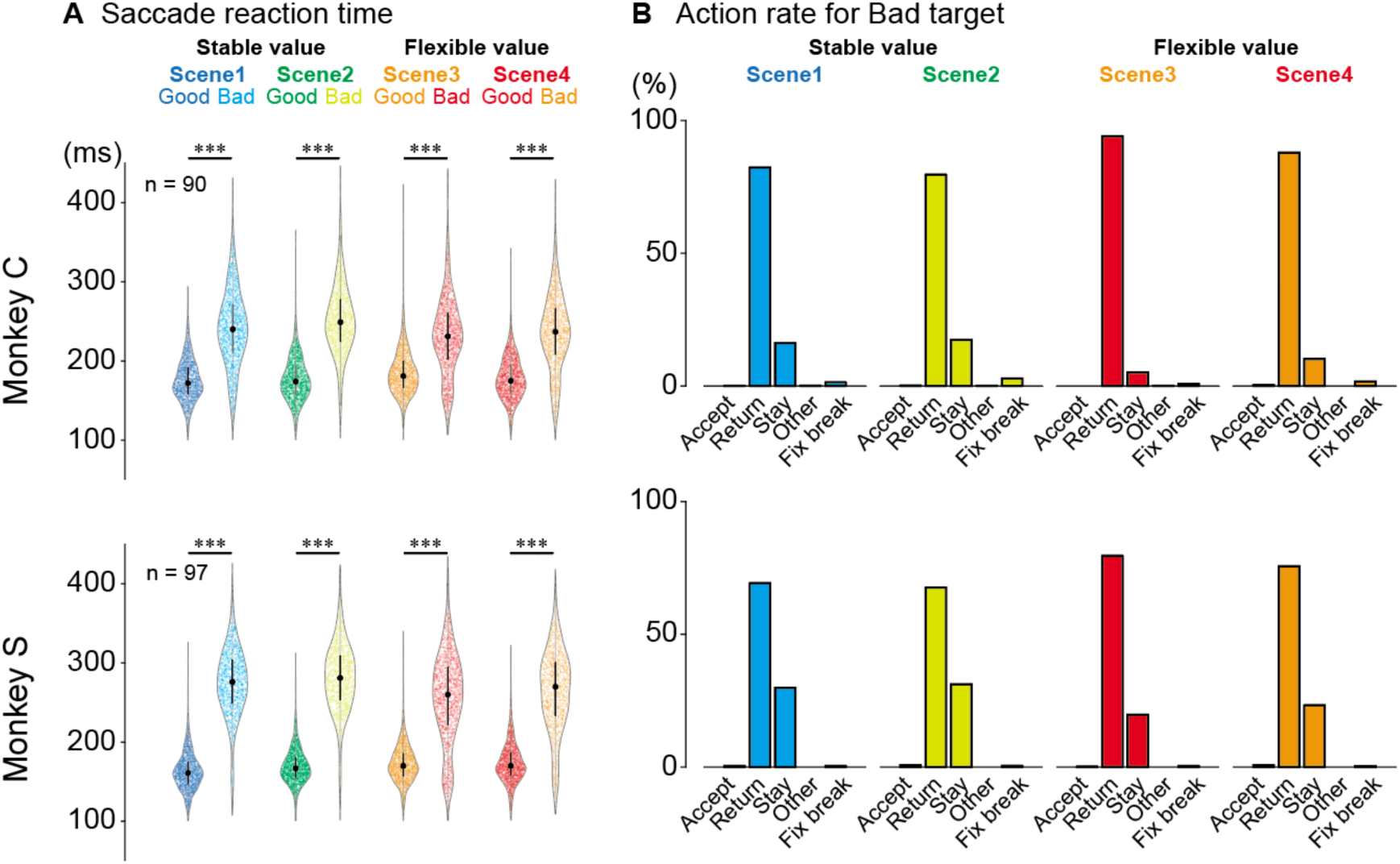
Behavioral performance. (A) Saccadic reaction times for good and bad objects in the two monkeys (C and S). Violin plots show that across all scenes and for both monkeys, saccades to good objects had significantly shorter latencies than saccades to bad objects during the return response (\*\*\**P* < 0.0001 for all comparisons, the Welch t-tests). Each dot represents data from a single recorded neuron (n = 90 for monkey C and n = 97 for monkey S). The thick horizontal line indicates the median, the box indicates the interquartile range (IQR), and the whiskers extend to the most extreme data point within 1.5 times the IQR. See also Table S1. (B) Proportions of different behavioral responses to bad objects for each scene. Both monkeys exhibited similar patterns: incorrect acceptance of bad objects was rare, and rejection responses consisted predominantly of return saccades, followed by stay and other responses. Accept incorrect acceptance of bad objects; Return, return saccades; Stay, maintaining central fixation; Other, saccades away from both the object and the central fixation point; Fix break, fixation break error. See also Table S2. Abbreviations: FP, fixation point; ITI, inter-trial interval; Obj, object; Tgt, target.

Consistent with our previous studies using this paradigm,^31,33,34^ the monkeys’ performance demonstrated a stable and robust understanding of the task rules and object values. Saccadic reaction times (RTs) were substantially faster for accepting good objects compared to rejecting bad objects via a return saccade (Figure S1A; Welch t-test, *P* < 0.0001). When faced with a bad object, the monkeys predominantly chose the “return” strategy over the “stay” strategy (Figure S1B). This behavioral pattern, which provides a reliable basis for analyzing neuronal activity related to action selection, was consistent across all recording sessions, and the same as in our previously published work.^31,33,34^

### General Firing Properties of STN Neurons

We recorded the activity of 187 task-responsive single neurons in the STN of two monkeys (Monkey C: 90; Monkey S: 97) as they performed the choice task. The subsequent analyses focused on trials with contralateral object presentations, which consistently elicited the most robust neuronal responses.

We observed a variety of firing patterns across our population of STN neurons in the sequential choice task. Figure 2 illustrates this variety with three representative single neurons. The most prominent and consistent responses across the population were sharp, phasic changes in firing rate immediately following target onset. However, the stimulus preferences in these responses varied across neurons. For example, some neurons increased their activity for both good and bad objects, with a preference for good objects (Figure 2A).

**Figure 2.**
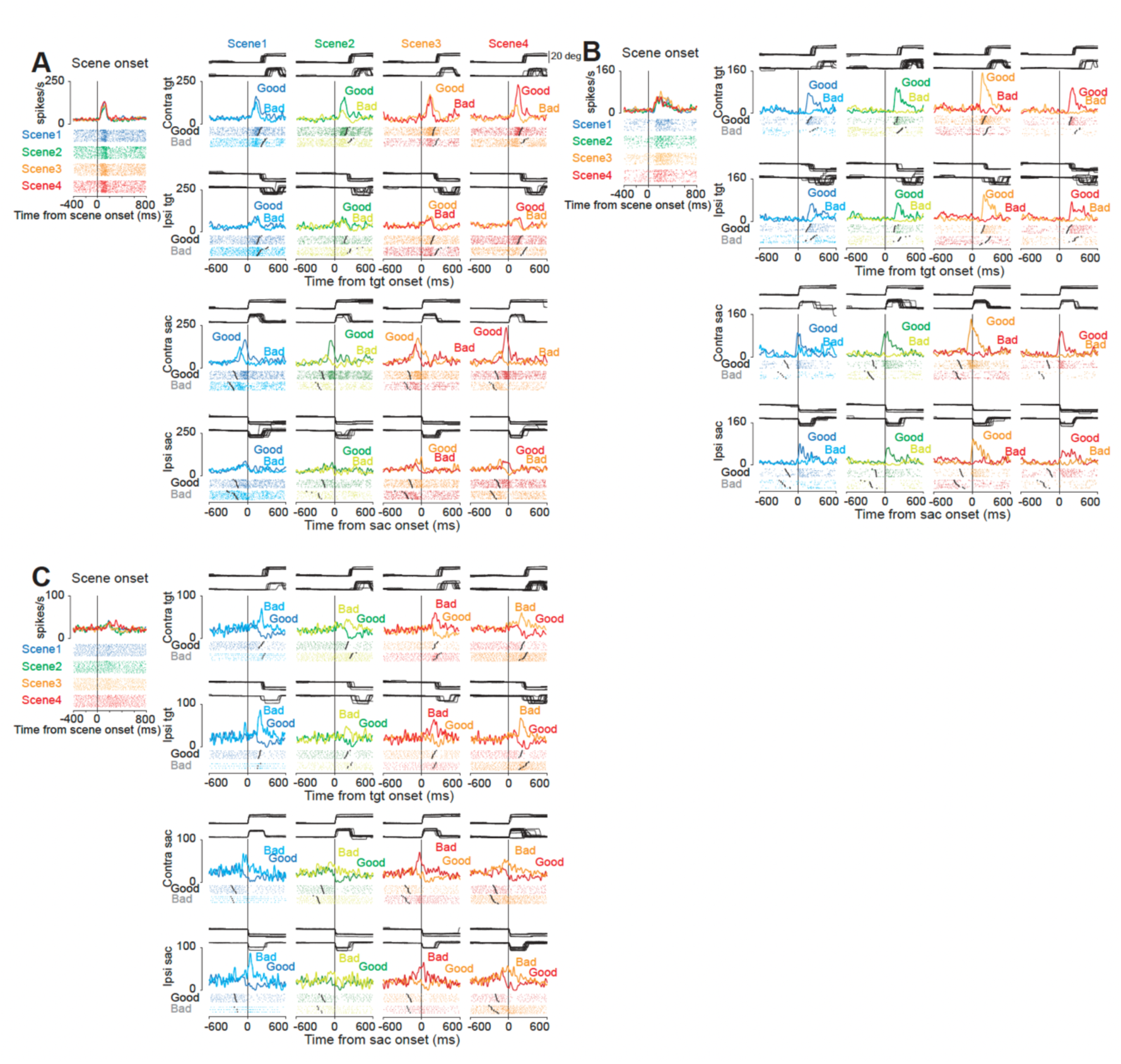
Representative neurons of the three clusters. (A, B, C) Activity of representative neurons from Cluster 1 (A), Cluster 2 (B), and Cluster 3 (C). For each neuron, the left columns show data aligned to scene onset or target onset, and the right columns show data aligned to saccade onset. Raster plots and spike density functions are shown for each condition. Colored traces indicate spike density for good and bad objects in each scene. In the target-aligned raster plots, larger circles indicate the time of saccade onset for each trial. In the saccade-aligned raster plots, larger circles indicate the time of target onset. Below each set of raster plots, horizontal eye position traces are shown, illustrating the timing and direction of saccades relative to each alignment event. Contralateral and ipsilateral target conditions are displayed separately.

Other neurons responded strongly to good objects but with little or no modulation for bad objects (Figure 2B). Finally, some neurons exhibited the opposite preferences and responded strongly to bad objects and were inhibited by good objects (Figure 2C).

This observed heterogeneity in firing patterns indicates that the STN is not a functionally uniform structure but instead contains distinct neuronal subpopulations, each potentially contributing differently to the decision-making process. To examine this possibility and objectively classify the neurons based on their activity profiles, we proceeded with a clustering analysis.

### Classification of STN Neurons into Functional Clusters

To objectively classify the diverse activity profiles observed in the STN, we used a k-means clustering algorithm based on the standardized firing rates during the presentation of contralateral good and bad objects (100–300 ms post-object onset). This specific methodology and time window were chosen to be identical to those used in our preceding study of the GPe, thereby allowing for a direct comparison with the functionally analogous neural populations we identified in the GPe.^31^ To quantitatively determine the optimal number of clusters (K), we simulated the silhouette values 5,000 times and found that K=3 produced the largest average silhouette value (Figure S2A). Accordingly, this data-driven analysis robustly partitioned the neuronal population into three distinct functional clusters (Figure S2B): Cluster 1 (n = 76), Cluster 2 (n = 90), and Cluster 3 (n = 21). We then examined whether these functionally defined groups also possessed distinct physiological or anatomical characteristics.

Comparison of spike waveform shapes revealed a significant difference in peak-to-peak duration across the three clusters (Kruskal-Wallis test, H = 8.39, *P =* 0.015; Figure S2C, D). Post-hoc pairwise comparisons with Bonferroni correction (α = 0.0167) showed that Cluster 1 and Cluster 3 differed significantly (*P* = 0.010), while Cluster 2 was intermediate (Cluster 1 vs Cluster 2: *P* = 0.071; Cluster 2 vs Cluster 3: *P* = 0.060). In contrast, baseline firing rates did not differ significantly among the three clusters (Kruskal-Wallis test, *P* > 0.05; Figure S2E).

**Figure S2.**
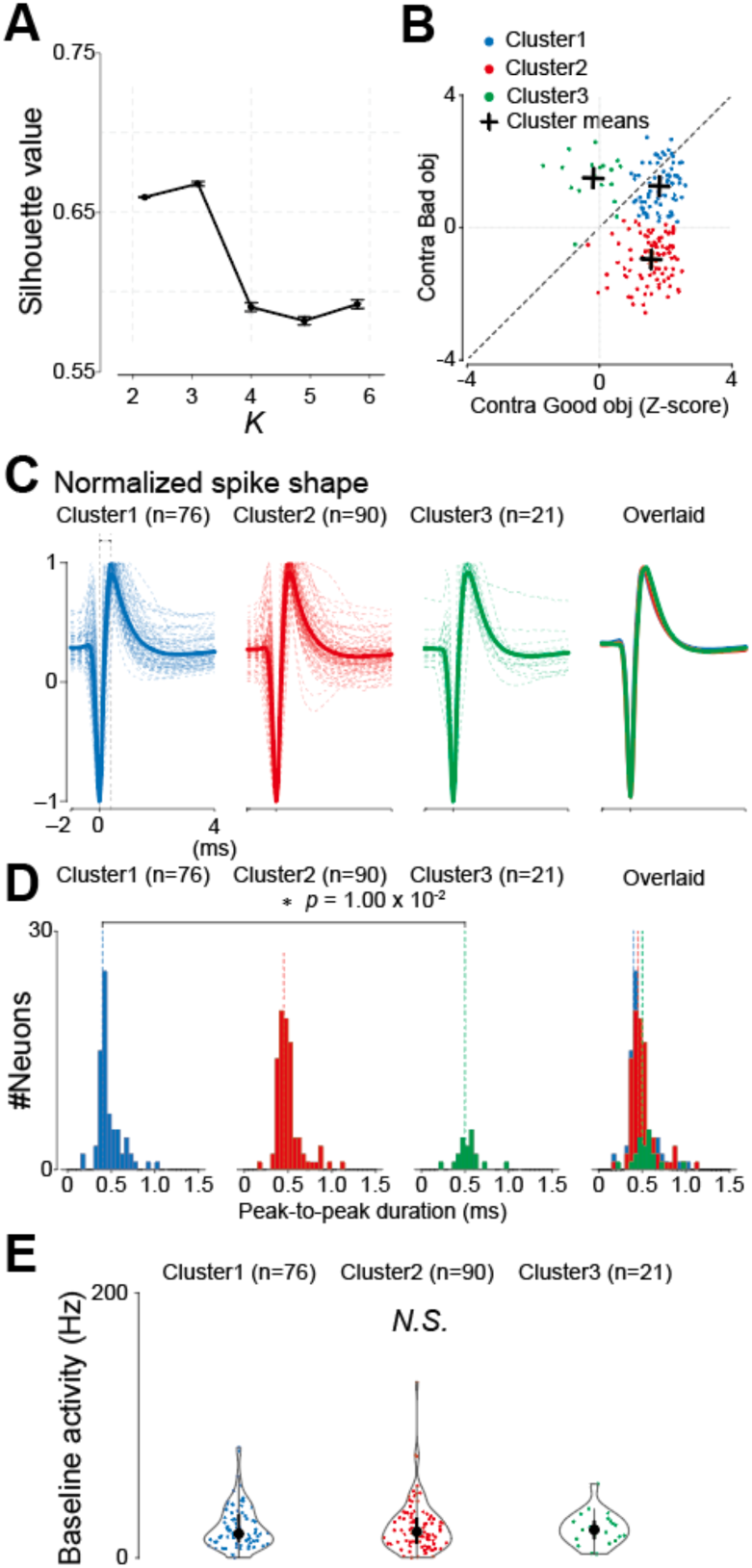
Classification and physiological properties of STN neuron clusters. (A) Optimal cluster number (K) determination. The plot shows the mean silhouette value (±SD) across 5,000 simulations for different values of K. K = 3 produced the largest average silhouette value. (B) Functional classification of STN neurons. Each point represents a single neuron, plotted according to its average z-scored firing rate in response to contralateral good (x-axis) and bad (y-axis) objects (100–300 ms post-object onset). Colors denote cluster assignment (blue: Cluster 1; red: Cluster 2; green: Cluster 3). Crosses indicate cluster centroids. (C) Normalized spike shapes for each cluster. The thick lines represent the average spike shape for each cluster, and the thin lines represent the individual spike shapes. Spike shapes were normalized to the trough and peak amplitudes for each neuron. (D) Distribution of peak-to-peak duration across clusters. Peak-to-peak duration differed significantly across clusters (Kruskal-Wallis test, H = 8.39, P = 0.015). Post-hoc pairwise comparisons with Bonferroni correction (α = 0.0167) revealed a significant difference between Cluster 1 and Cluster 3 (P = 0.010), while other pairs did not reach significance (Cluster 1 vs Cluster 2: P = 0.071; Cluster 2 vs Cluster 3: P = 0.060). (E) Baseline firing rates for each cluster. No significant difference was found among the clusters (Kruskal-Wallis test, P > 0.05). Plot conventions for violin plots (D, E): circle, median; thick line, interquartile range (IQR); thin line, 1.5 × IQR.

### Recording site distribution

Recording electrodes were advanced vertically from a dorsal approach through a recording grid mounted on the implanted chamber. Representative MRI sections showing the electrode trajectories and STN locations for each monkey are presented in Figure S3A. Three-dimensional reconstruction of all recording sites confirmed that neurons across all three clusters were concentrated in the ventral portion of the STN in both monkeys (Figure S3B). One-sample t-tests comparing cluster coordinates against the STN centroid revealed a consistent ventral bias across multiple cluster-hemisphere combinations (*P* < 0.05). To test whether the three functional clusters occupied distinct subregions within the STN, we performed Kruskal-Wallis tests comparing coordinates across clusters for each axis, hemisphere, and monkey. No significant topographic segregation was found after FDR correction (all corrected P > 0.06), indicating that neurons with different functional properties were spatially intermingled within the ventral STN.

**Figure S3.**
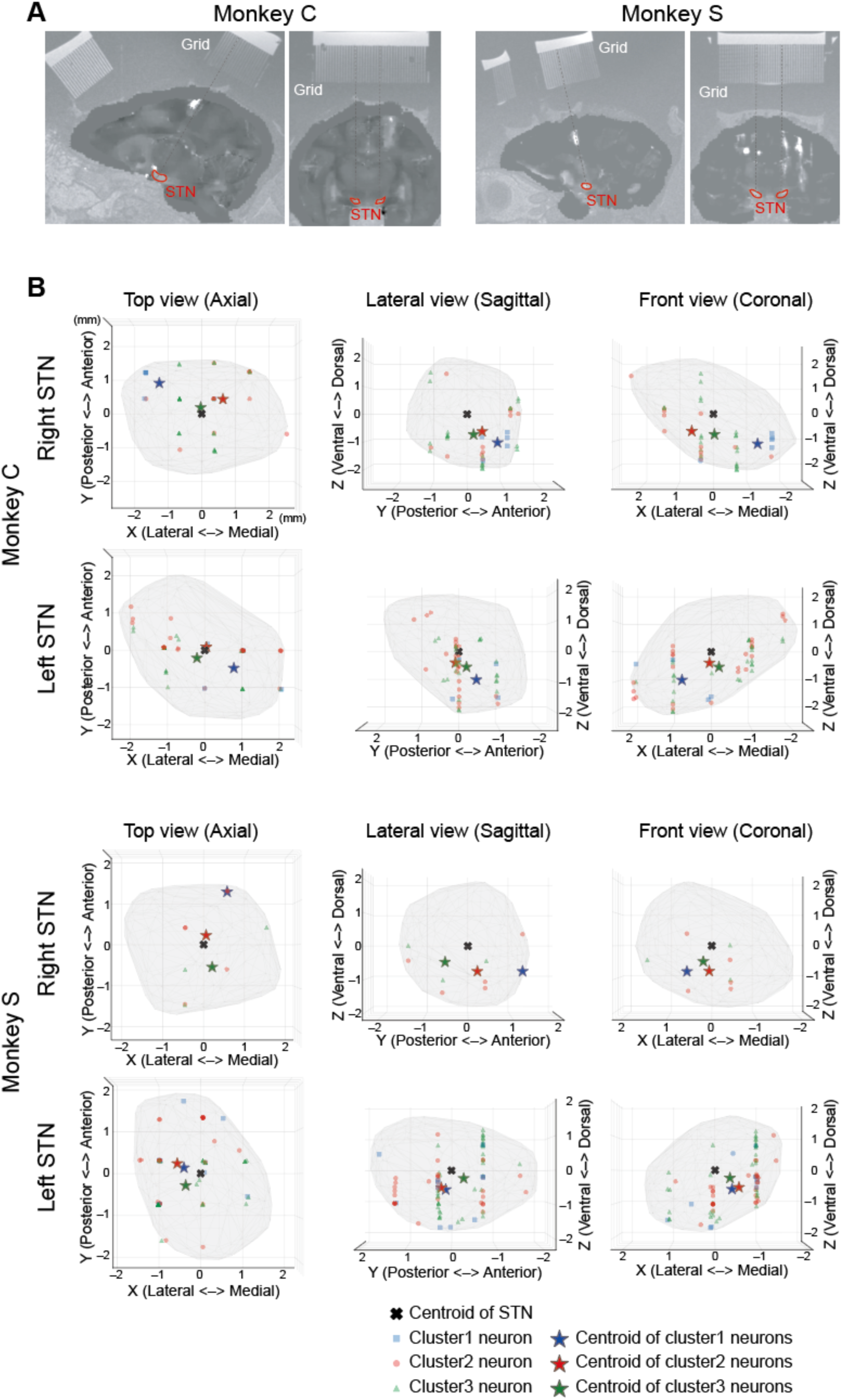
Electrode trajectories and anatomical distribution of recording sites within the STN. (A) Representative MRI sections showing electrode trajectories and STN locations for Monkey C (left) and Monkey S (right). Images were reconstructed by merging high-resolution grid scan images, which visualize the recording grid holes used for electrode guidance, with quantitative susceptibility mapping (QSM) images that delineate the STN (see Methods for details). Sagittal (left) and coronal (right) sections are shown for each monkey. Dashed lines indicate the electrode insertion trajectories through the recording grid. Electrodes were inserted vertically through a recording grid mounted on the chamber, approaching the STN from a dorsal direction. (B) Recording locations of all task-responsive neurons shown for each monkey and hemisphere in three orthogonal projections: axial (top view, left column), sagittal (lateral view, middle column), and coronal (front view, right column). Coordinates are expressed in millimeters relative to the STN centroid (origin), computed as the mean of all STN boundary vertices for each hemisphere. Small symbols indicate individual neurons (blue squares: Cluster 1; red circles: Cluster 2; green triangles: Cluster 3). Large star symbols indicate the centroid of recording locations for each cluster (blue: Cluster 1; red: Cluster 2; green: Cluster 3). The dark cross indicates the STN centroid. Gray shading represents the STN boundary derived from template-based segmentation validated against quantitative susceptibility mapping (see Methods). Recordings across all three clusters were concentrated in the ventral part of the STN. No significant topographic segregation among clusters was detected after FDR correction (Kruskal-Wallis tests).

### Task-Related Activity Revealed Distinct Value and Spatial Coding

To dissect the functional roles of these clusters, we examined their population activity aligned to scene onset and object onset (Figure 3). As would be expected from our clustering method, the activity of the neurons in each cluster was clearly different following the presentation of the target objects. In contrast, none of the clusters showed significant modulation related to the scene context itself (Figure 3A, B, E, F, I, J). We therefore characterized the specific nature of this target-evoked activity for each of the three clusters.

**Figure 3.**
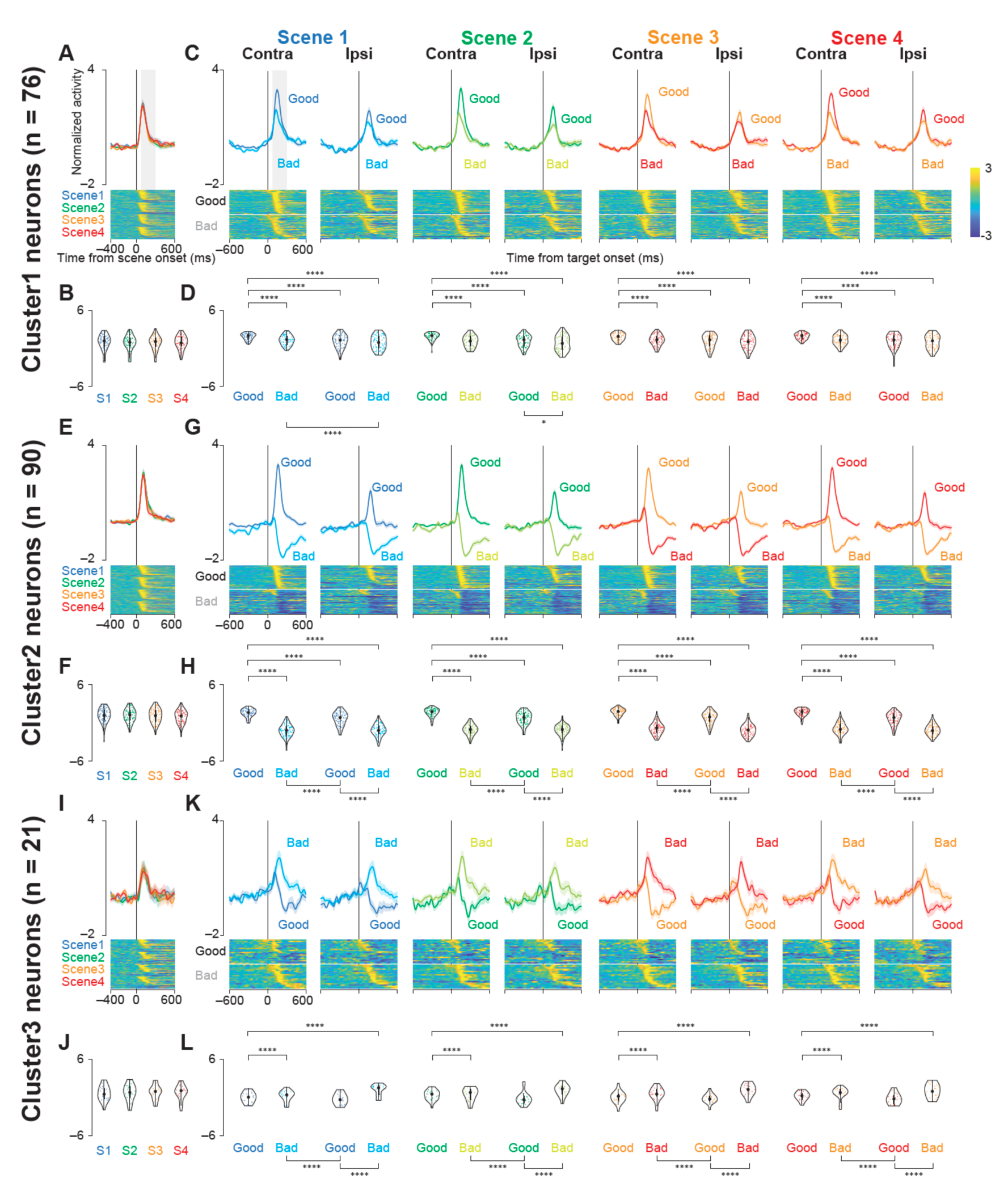
Population activity of three groups of STN neurons at scene and target onsets during the choice task. (A, E, I) Mean normalized population activity (activity was normalized by subtracting the baseline firing rate and dividing it by the standard deviation) of Cluster 1 (A), Cluster 2 (E), and Cluster 3 (I) neurons aligned to the onset of the scene (scenes 1–4) during the choice task. The shaded areas indicate mean ± standard errors of mean (SEMs). The lower panels show color maps of the normalized activity of individual neurons. Each row in the color map represents a single neuron, and the neurons are sorted based on the time at which their standardized activity exceeds a threshold. (B, F, J) Violin plots showing the distribution of mean normalized neuronal activity of individual neurons in Clusters 1 (B), 2 (F), and 3 (J) for each scene onset during the choice task. Neuronal activity was measured at 200-ms intervals beginning 100 ms after scene onset. The larger circle indicates the median value, the thick vertical line shows the interquartile range (IQR), and the thin vertical line indicates 1.5 × IQR. (C, G, K) Mean normalized population activity of Cluster 1 (C), Cluster 2 (G), and Cluster 3 (K) neurons aligned to the onset of contralateral and ipsilateral good and bad targets for scenes 1-4. (D, H, L) Violin plots showing the distribution of the mean normalized neuronal activity of individual neurons in Clusters 1 (D), 2 (H), and 3 (L) for each target-onset condition. Neuronal activity was measured for a 200-ms interval beginning 100 ms after target onset. The asterisks indicate significant differences in neuronal activity among the four conditions (contra good, contra bad, ipsi good, ipsi bad) within each scene (post hoc pairwise t tests with Bonferroni correction, *P* < 0.05, etc.). See also Tables S3–S5.

The activity of Cluster 1 neurons increased following the presentation of both good and bad objects, though the response was significantly stronger for good objects (Figure 3C, D; *P* < 0.001, see Table S3 for statistics). These neurons also showed a clear preference for contralateral targets, indicating that they integrated both value and spatial information.

Cluster 2 exhibited a distinct, bidirectional response profile. Upon target presentation, these neurons increased their firing for good objects but decreased their firing for bad objects (Figure 3G, H; *P* < 0.001, Table S4). This opponent coding of positive and negative value, with a strong response to contralateral good targets, points to a primary role for this cluster in value-based action selection.

In contrast, Cluster 3 appeared to preferentially encode negative outcomes. While their activity increased for both object types, the response was significantly more pronounced for bad objects compared to good objects (Figure 3K, L; *P* < 0.001, Table S5). This response pattern is consistent with a role in encoding the negative value of presented objects, a functional interpretation that is further examined below with temporal alignment and response strategy analysis.

Together, these findings demonstrate that the STN contains functionally specialized subpopulations, each processing a unique combination of value and spatial information to guide behavior.

### Validation of Functional Groupings with a Data-Driven PCA Approach

To validate the functional groupings identified by our window-based method and to explore the underlying structure of the data in a more objective, data-driven manner, we next employed Principal Component Analysis (PCA). To focus on the evoked responses that are central to this study, we performed this analysis on the standardized neural activity from a short window of 0 to 300 ms after the onset of contralateral good and bad objects.

This analysis revealed the principal temporal patterns (components) that explain the variance across the neural population (Figure S4A), with a Scree plot indicating that most of the variance was captured by the first few components (Figure S4B). We then performed k-means clustering on the weights of each neuron on the top three PCs (PC1-3), which can be visualized by considering the position of each neuron in a 3-dimensional functional space (Figure S4C).

To fully characterize the possible functional groupings, we examined the solutions for both K=2 and K=3. The results of this analysis are shown in Figure S4, which displays the population activity and the normalized neuronal activity of individual neurons.

Both clustering solutions consistently support our primary findings. The K=2 solution revealed a fundamental division of the neurons into a ‘value-coding facilitative’ group and a larger ‘general facilitative’ group. When the number of clusters was increased to K=3, the analysis largely preserved the distinct ‘value-coding’ group, while subdividing the ‘general facilitative’ group into two smaller sub-groups.

Taken together, this data-driven analysis is consistent with our main conclusion that the activity of STN neurons in our task is organized around principles of action facilitation. The core distinction between value-coding and general facilitation was observed regardless of the clustering details, lending further support to our claim that the role of the STN in this context is largely facilitative. Notably, a functionally distinct negative value-coding group, analogous to Cluster 3 from our initial analysis, was not robustly isolated by this method, a point we will consider further in the Discussion.

**Figure S4.**
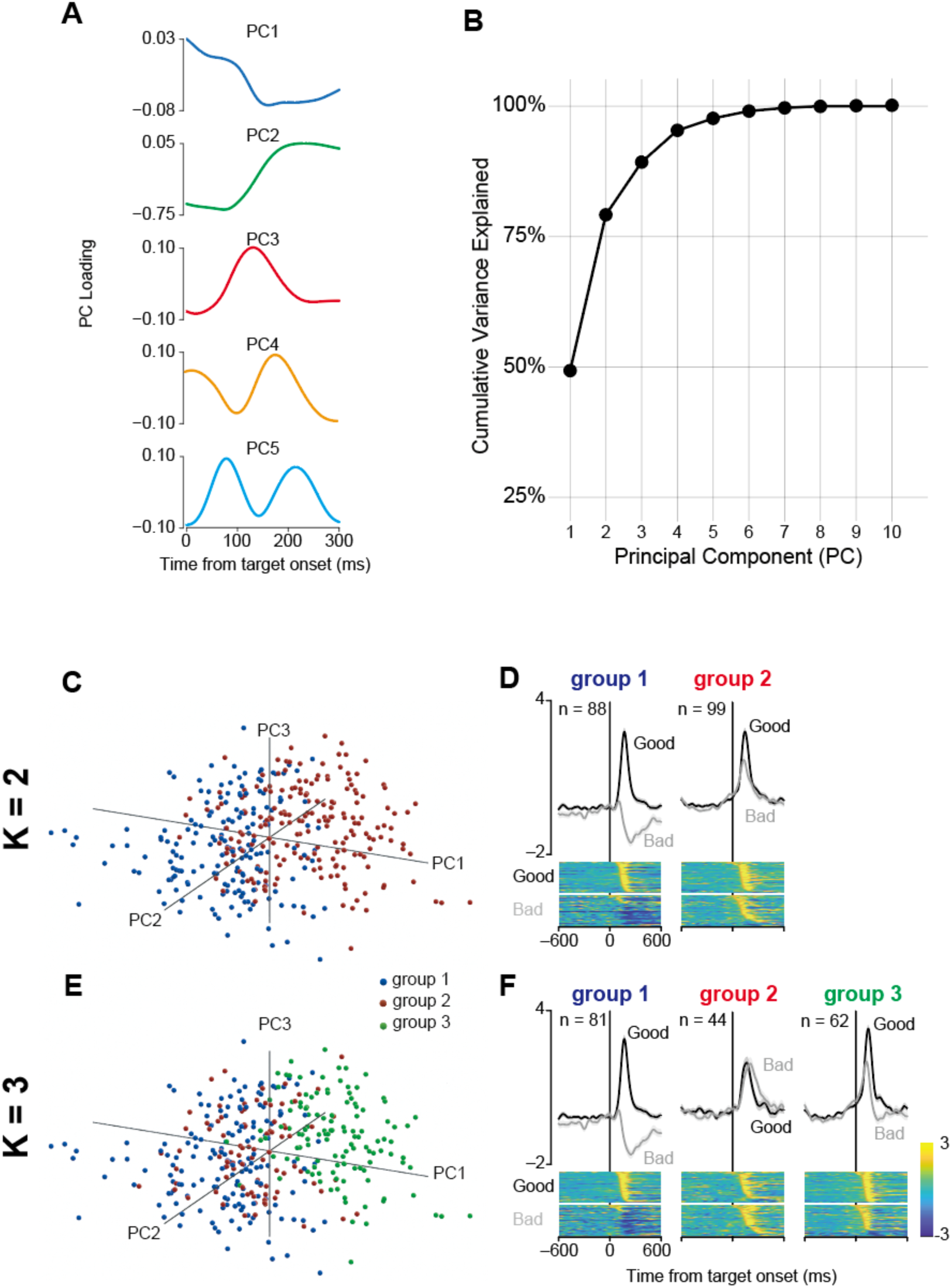
PCA-based functional grouping of STN neurons using post-stimulus activity. (A) Temporal waveforms of the top five principal components (PCs) derived from the neural activity in the 0–300 ms post-stimulus window. The x-axis represents time from target onset, and the y-axis represents the PC loading. (B) Cumulative variance plot. The curve shows the cumulative percentage of total variance explained as a function of the number of principal components included. (C) 3D scatter plot of the neural population in the space defined by the top three PCs. Each point represents a single neuron-condition, colored by its cluster assignment for the K=2 solution. (D) Population activity for the two functional groups identified at K=2. Top panels show the mean normalized activity (± SEM) for contralateral good (black) and bad (gray) trials. Bottom panels show heatmaps of the Z-scored activity of all individual neurons within each group, sorted by response latency. (E) Same as (C), but for the K=3 clustering solution. (F) Same as (D), but for the three functional groups identified at K=3.

### Saccade-Related Activity Reflected Action Choice

To simplify the analysis of movement-related activity, we focused on data from Scene 1, as target-aligned responses were consistent across all scenes. To investigate the relationship between neuronal activity and movement execution, we next aligned the firing of each cluster to saccade onset (Figure 4).

**Figure 4.**
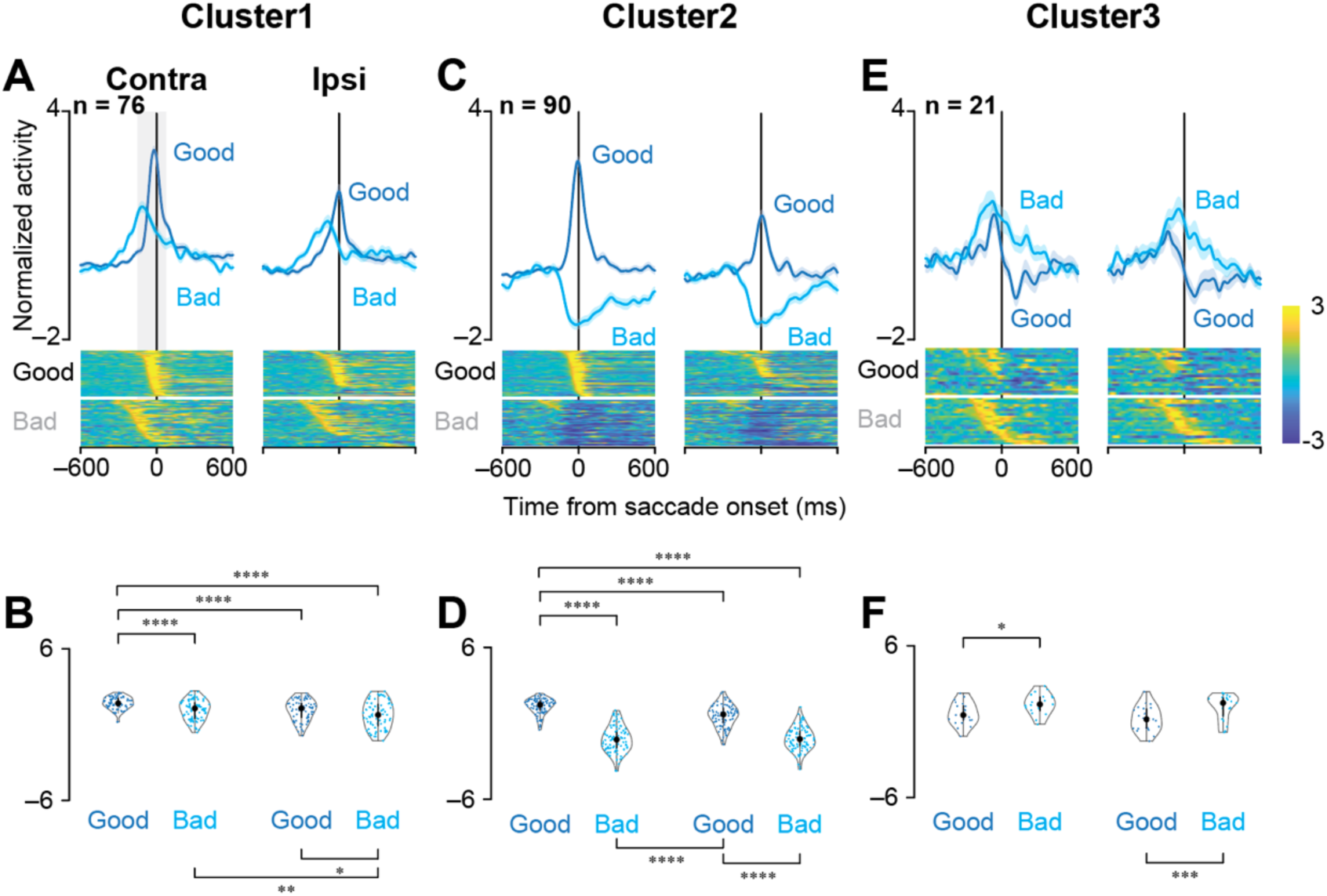
The population activity of the three groups of STN neurons aligned with saccade onset during the choice task. Note that because the reaction times for rejecting “bad” objects were longer than those for accepting “good” objects (see Figure S1A), stimulus-related activity for “bad” trials appears shifted earlier in time relative to saccade onset. This figure illustrates activity relative to the motor event, not the stimulus-evoked response latency. (A, C, E) Mean normalized population activity of Cluster 1 (A), Cluster 2 (C), and Cluster 3 (E) neurons aligned to saccade onset for contralateral or ipsilateral, and good or bad objects in Scene 1 during the choice task. The shaded areas indicate ± SEMs. The lower panels show color maps of normalized neuronal activity of individual neurons, with each row representing a single neuron, sorted as in Figure 3. (B, D, F) Violin plots showing the distribution of the mean normalized neuronal activity of individual neurons in Clusters 1 (B), 2 (D), and 3 (F) when monkeys made a saccade to the target. Neuronal activity was measured at 200-ms intervals from 150 ms before to 50 ms after saccade onset. The asterisks indicate significant differences in neuronal activity among the four conditions (*P* < 0.05, etc.; post hoc pairwise t tests with Bonferroni correction). See also Tables S6–S8.

The activity of neurons in Cluster 1 was significantly elevated around the time of saccade initiation, with the highest firing rates observed for saccades to contralateral good objects (Figure 4A, B; *P* < 0.001, see Table S6 for statistics). This robust peri-movement activity suggests a close association with the generation of saccades.

Neurons in Cluster 2 also showed strong modulation at saccade onset, but their activity was again contingent on value. Firing increased for saccades to good objects and decreased for those to bad objects (Figure 4C, D; *P* < 0.001, Table S7), reinforcing the idea that neurons in this cluster integrate value and spatial signals in relation to saccade execution.

In contrast, neurons in Cluster 3 showed the greatest increase in activity preceding saccades to bad objects (Figure 4E, F; *P* < 0.001, Table S8). This response pattern is opposite to that of the facilitative clusters and is consistent with encoding the negative value of the presented stimulus. However, because the reaction times for bad objects were longer than those for good objects (Figure S1A), the apparent pre-saccadic activity in this alignment could also reflect a stimulus-locked response that appears early relative to the delayed saccade. The temporal alignment analysis below directly addresses this ambiguity.

### Temporal alignment heterogeneity across STN clusters

To determine whether STN neuronal responses were time-locked to the sensory event (target onset) or to the motor response (saccade onset), we computed a Locking Index for each neuron based on the trial-by-trial variability of response onset latency when aligned to target onset (SD_target) versus saccade onset (SD_saccade), adapting the framework of the previous study.^35^ Single-trial onsets were detected within the early post-target window (100–300 ms; see Methods), so this analysis specifically probes alignment of the early task-evoked response component. The Locking Index was defined as (SD_target − SD_saccade) / (SD_target + SD_saccade), where negative values indicate target-locked activity and positive values indicate saccade-locked activity (Figure 5A). The majority of neurons across all clusters were target-locked: 68% (52/76) in Cluster 1, 76% (68/90) in Cluster 2, and 85% (18/21) in Cluster 3 (Figure 5B).

**Figure 5.**
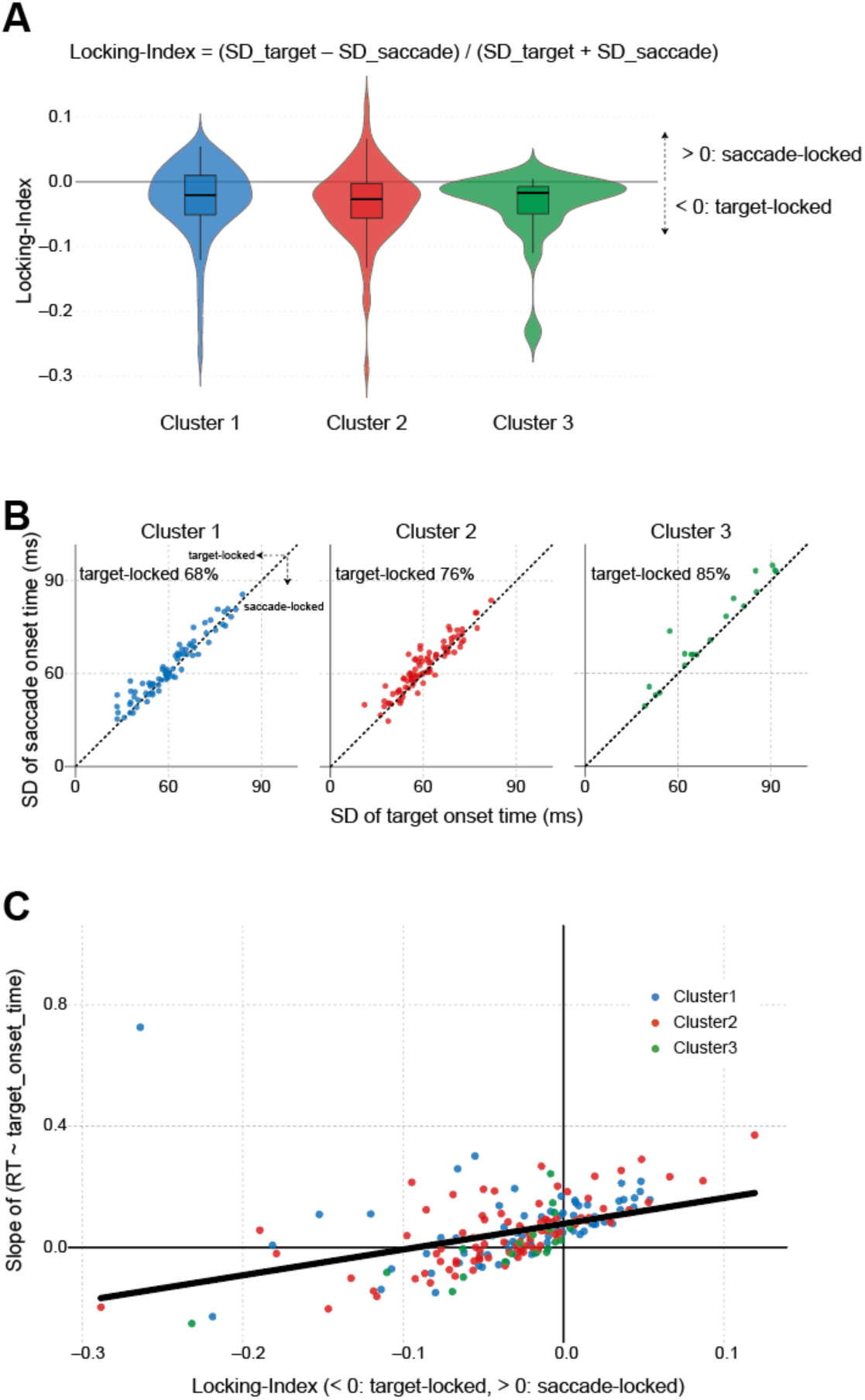
Temporal alignment properties of STN neurons. (A) Distribution of Locking Index across clusters. The Locking Index was defined as (SD_target − SD_saccade) / (SD_target + SD_saccade), where SD_target and SD_saccade represent the standard deviation of single-trial response onset latencies measured relative to target onset and saccade onset, respectively (see Methods). Negative values indicate target-locked activity; positive values indicate saccade-locked activity. Box plots within the violins show the median and interquartile range. (B) Relationship between SD of response onset latency relative to target onset (x-axis) and relative to saccade onset (y-axis) for each neuron. The dashed diagonal line indicates equal variability (SD_target = SD_saccade); neurons above this line (SD_target < SD_saccade) are target-locked, whereas neurons below this line (SD_target > SD_saccade) are saccade-locked. The percentage of target-locked neurons is indicated for each cluster (Cluster 1: 68%, Cluster 2: 76%, Cluster 3: 95%). (C) Relationship between Locking Index and the slope of a per-neuron linear regression of saccadic reaction time on response onset latency (measured relative to target onset). A slope near 0 is expected for target-locked neurons, whose onset timing is independent of RT; a slope approaching 1 is expected for saccade-locked neurons, whose onset timing covaries with saccade initiation. The solid line shows the linear fit across all neurons. Each dot represents one neuron; colors indicate cluster identity (blue: Cluster 1; red: Cluster 2; green: Cluster 3).

To validate this classification with a complementary approach, we investigated whether the response onset latency of each neuron (relative to target onset) predicted saccadic reaction time on a trial-by-trial basis. For each neuron, we regressed RT on response onset latency and extracted the slope. A slope near 0 is expected for target-locked neurons, whose onset timing is independent of RT; a slope near 1 is expected for saccade-locked neurons, whose onset timing covaries with saccade initiation. The Locking Index and slope were strongly correlated across all neurons (Spearman ρ = 0.631, *P* < 2.2 × 10⁻¹⁶; Figure 5C), confirming that these two independent measures converge. This correlation was consistent within each cluster (Cluster 1: ρ = 0.563, *P* = 2.1 × 10⁻⁷; Cluster 2: ρ = 0.680, *P* < 10⁻¹⁰; Cluster 3: ρ = 0.833, *P* = 5.5 × 10⁻⁷).

### Spatial and value coding properties across clusters

To characterize the spatial selectivity of each cluster, we examined the contralateral-ipsilateral difference for good and bad objects separately using post-hoc comparisons from the Linear Mixed-effects Models (LMMs) (Tables S3–S5). The three clusters exhibited distinct profiles of spatial coding. Cluster 1 showed a significant contralateral preference for both good objects (all scenes *P* < 0.0001) and bad objects (all scenes *P* < 0.012), indicating that this population integrates both value and spatial information. Cluster 2 exhibited a strong contralateral preference exclusively for good objects (all scenes *P* < 0.0001), while the contralateral-ipsilateral difference was absent for bad objects (*P* > 0.05 in most scenes). This asymmetry is consistent with the bidirectional value coding of Cluster 2, in which activity is suppressed for bad objects and the spatial signal is consequently attenuated. In contrast, Cluster 3 showed no significant contralateral preference for either good or bad objects (all *P* > 0.06), indicating a non-directional response.

Together with the temporal alignment analysis, these results reveal a systematic functional organization across the three STN clusters. Clusters 1 and 2, which comprise the facilitative majority, are predominantly target-locked with strong spatial selectivity, consistent with stimulus-driven visuomotor signals modulated by object value. The saccade-locked minority within these clusters may correspond to a more motor-related processing stage. Cluster 3, by contrast, is almost exclusively target-locked and non-directional, consistent with a pure value signal that is not mapped to a specific motor output.

### Activity in Facilitative Clusters Was Associated with Reaction Time

The preceding analyses suggest that neurons in clusters 1 and 2 are involved in facilitating saccades. If this is the case, then trial-to-trial fluctuations in their neuronal activity should be associated with corresponding variations in reaction time (RT). To examine this, we analyzed the relationship between RT and three neuronal response parameters (onset latency, peak magnitude, and slope) using LMMs (Figure S5, also see Table S9 for full model results). This approach allowed us to account for non-independent observations from the same monkeys and neurons. To correct for multiple comparisons across the nine models fitted, we applied a Bonferroni correction, adopting a significance threshold of *P* < 0.0056 (0.05/9).

After applying this correction, the LMM analysis indicated that only the response onset latency in the facilitative clusters had a statistically significant, albeit modest, association with RT. The population-level relationship between response onset latency and saccadic reaction time was statistically significant but modest in magnitude (Cluster 1: β = 0.036, *P* = 0.0004; Cluster 2: β = 0.057, *P* < 0.0001). This modest association is consistent with the temporal alignment analysis described above. Because most neurons were target-locked, their onset timing did not vary with RT, and this weakened the overall relationship.

The significant association was driven by the saccade-locked minority, whose onset timing varied together with saccade initiation. Thus, the small beta coefficients do not reflect a uniformly weak relationship across the population but rather arise from mixing two groups of neurons with different temporal properties within each cluster.

In contrast, neither the peak firing magnitude nor the response slope for these facilitative clusters showed a significant relationship with RT after Bonferroni correction (all *P* > 0.0056). Furthermore, consistent with the uncorrected analysis, no activity parameters from Cluster 3 significantly predicted RT.

Taken together, these results suggest a subtle link between the trial-by-trial activity in the facilitative STN populations and the speed of action initiation, whereas no such link was found for the negative value-coding population (Cluster 3). Notably, after rigorous statistical correction, this weak association appears to be specific to the timing of the neuronal response, as only the onset latency, and not the response magnitude or slope, remained a significant predictor of behavior.

**Figure S5.**
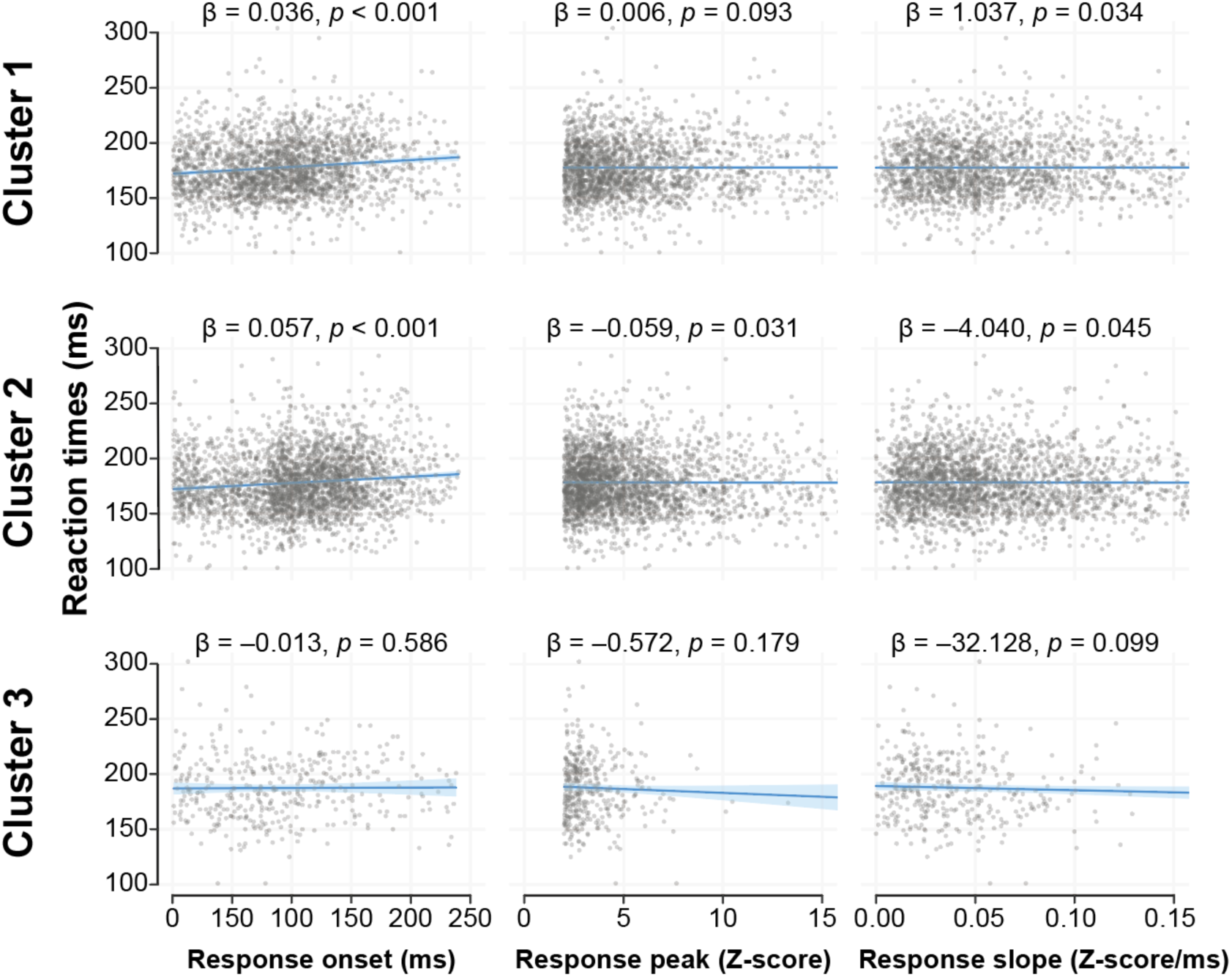
Trial-by-trial relationship between STN neuronal activity and reaction times. Scatter plots showing the relationship between saccadic RTs and three neuronal response parameters for each STN cluster. Each gray dot represents a single trial under the ‘good, contra, accept’ condition. Rows and columns correspond to the three neuronal clusters and three response parameters (onset latency, peak magnitude, and slope), respectively. Lines and shaded ribbons represent the predicted marginal effect and 95% confidence interval, respectively, derived from the LMM (linear mixed-effects model). Inset values indicate the unstandardized regression coefficient (β) and the corresponding *P*-value from the LMM. *P*-values shown are uncorrected; interpretation in the main text is based on a Bonferroni-corrected significance threshold (*P* < 0.0056). Full model results are provided in Table S9.

### Cluster 3 activity was independent of rejection strategy

In our task, monkeys rejected bad objects using two main strategies, return saccades (redirecting gaze back to the fixation point) and stay responses (maintaining central fixation without making a saccade). If Cluster 3 functions as a motor suppression signal, its activity should differ between these strategies, as they involve different motor outputs. To test this, we compared Cluster 3 population activity between return and stay trials for bad objects using an LMM. No significant difference was found for either contralateral (*P* = 0.512) or ipsilateral (*P* = 0.121) targets (Figure S6). This analysis was restricted to choice task trials, as the fixation task did not involve a choice between response strategies. These results demonstrate that Cluster 3 activity reflects the value of the presented object rather than the motor strategy used to respond to it.

**Figure S6.**
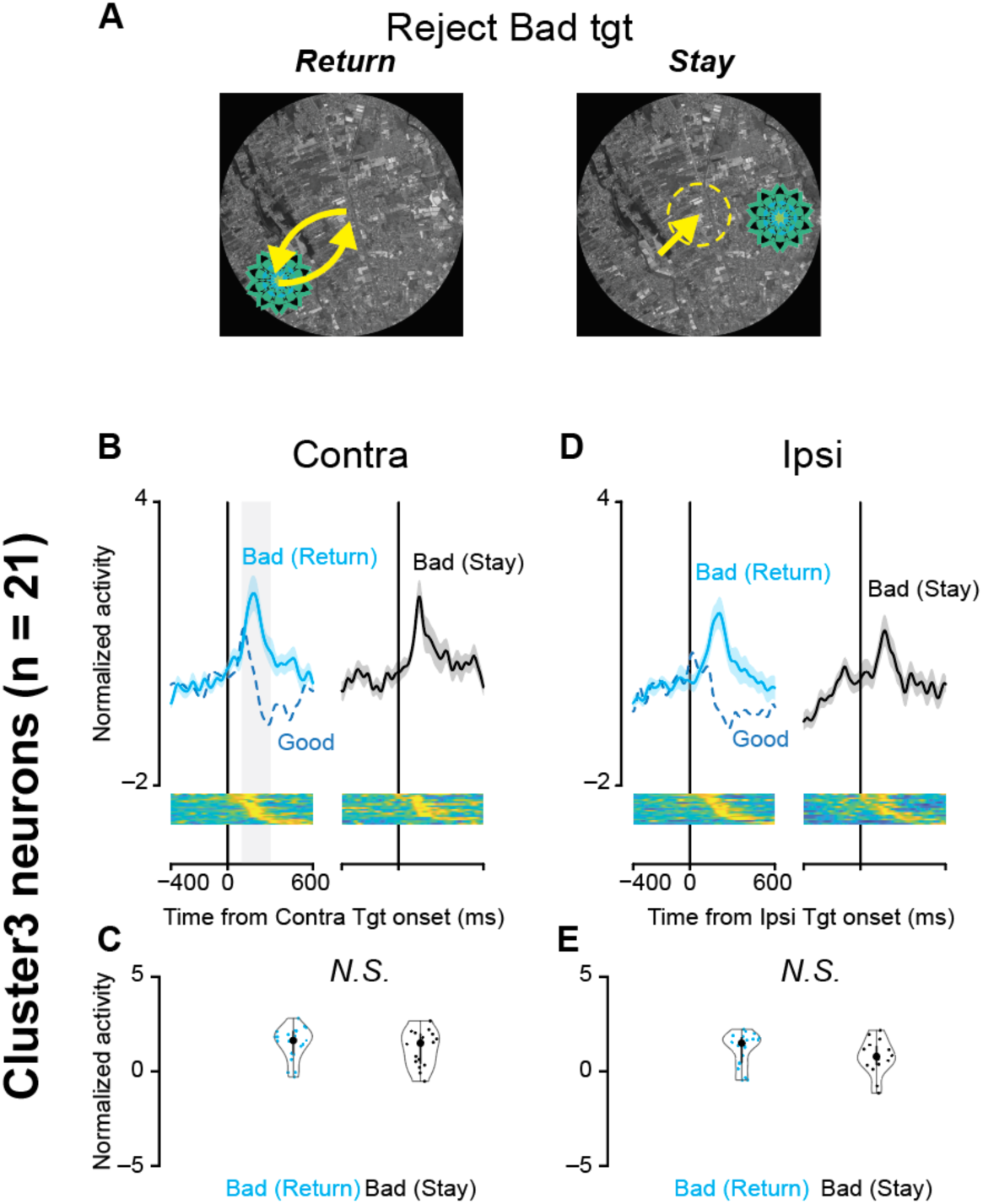
Cluster 3 activity was independent of rejection strategy. (A) Schematic illustration of the two rejection strategies for bad objects. When a bad object appeared as a target, monkeys could reject it either by making a saccade back to a previously visited location (Return) or by maintaining fixation (Stay). (B) Population spike density functions of Cluster 3 neurons (n = 21) aligned to contralateral target onset for bad objects rejected by return (cyan) and bad objects rejected by stay (black), shown together with the response to good objects (blue dot) for reference. Shaded region indicates the analysis window (100–300 ms after target onset). (C) Violin plots comparing mean normalized activity during the 100–300 ms post-target window between return and stay conditions for contralateral bad objects. Each symbol represents one neuron. No significant difference was found (LMM, *P* = 0.512). (D, E) Same as (B, C) but for ipsilateral target presentations. No significant difference was found between return and stay conditions (*P* = 0.121). These results indicate that Cluster 3 activity reflects the value of the presented object rather than the specific motor strategy used to reject it.

### Response latency hierarchy across STN clusters

To examine whether the three functional clusters differ in the speed of their responses, we compared neuronal response latencies across clusters (Figure S7). Response latency was defined as the first time point (100–300 ms post-event) at which z-scored activity exceeded 2.0 for at least 5 consecutive milliseconds. For target onset conditions, latency was measured for the preferred stimulus of each cluster (good objects for Clusters 1 and 2; bad objects for Cluster 3) to ensure reliable onset detection.

At scene onset, Clusters 1 and 2 responded with similar latencies (*P* = 6.94 × 10⁻², N.S. after Bonferroni correction), while both responded significantly earlier than Cluster 3 (Cluster 1 vs Cluster 3: *P* = 4.17 × 10⁻⁵; Cluster 2 vs Cluster 3: *P* = 1.55 × 10⁻⁵). Following contralateral target onset, Cluster 1 responded earliest (Cluster 1 vs Cluster 2: *P* = 3.11 × 10⁻¹³; Cluster 1 vs Cluster 3: P = 5.47 × 10⁻⁵), while Clusters 2 and 3 did not differ significantly (*P* = 3.91 × 10⁻², N.S. after Bonferroni correction). A similar pattern was observed for ipsilateral target onset (Cluster 1 vs Cluster 2: *P* = 3.23 × 10⁻²³; Cluster 1 vs Cluster 3: *P* = 5.44 × 10⁻⁶; Cluster 2 vs Cluster 3: *P* = 7.69 × 10⁻¹). All pairwise comparisons used two-sample Kolmogorov-Smirnov tests with Bonferroni correction (α = 0.05/3 = 0.0167).

These results reveal a latency hierarchy across the three clusters. Cluster 1 provides the earliest response, consistent with a rapid visuospatial signal. Cluster 2 responds at intermediate latencies, suggesting additional processing for value discrimination. Cluster 3 responds latest at scene onset, consistent with a more evaluative role in assessing the negative value of presented objects.

**Figure S7.**
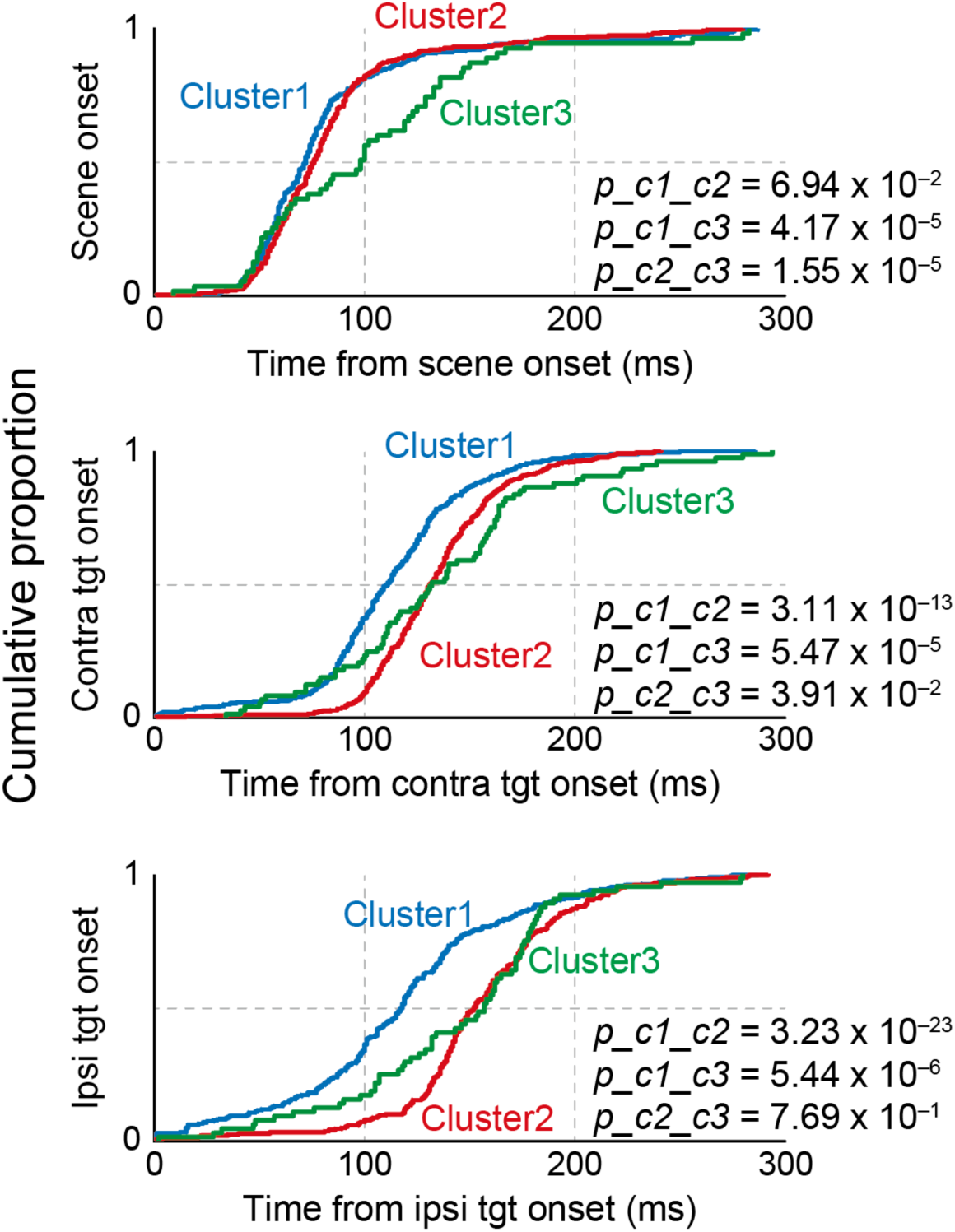
Comparison of response latencies across STN clusters. Cumulative distribution functions (CDFs) of neuronal response latency for Cluster 1 (blue), Cluster 2 (red), and Cluster 3 (green), aligned to scene onset (top), contralateral target onset (middle), and ipsilateral target onset (bottom). Response latency was defined as the first time point (100–300 ms post-event) at which z-scored activity exceeded 2.0 for at least 5 consecutive milliseconds (see Methods). For scene onset, all clusters were compared on the identical event. For target onset, latency was measured for the preferred stimulus of each cluster (good objects for Clusters 1 and 2; bad objects for Cluster 3) to ensure reliable onset detection. Data were pooled across scenes 1–4, as individual neurons showed consistent response patterns across scenes. *P*-values are from two-sample Kolmogorov-Smirnov tests (Bonferroni-corrected significance threshold: α = 0.05/3 = 0.0167). At scene onset, Clusters 1 and 2 responded with similar latencies, while both preceded Cluster 3. Following target onset, Cluster 1 exhibited the shortest latencies, responding significantly earlier than both Cluster 2 and Cluster 3 in both directions.

### STN response latencies preceded those of functionally analogous GPe populations

To test the hypothesis that STN activity exerts a feedforward excitatory effect on the GPe, we compared neuronal response latencies between STN clusters and functionally analogous GPe populations recorded during the identical paradigm in the same monkeys.^31^ STN Cluster 1 was compared with GPe Cluster 1, and STN Cluster 2 was compared with GPe Cluster 2, based on their analogous response profiles (Figure 6A). Response latencies were measured using the same criteria as described above (z-score threshold of 2.0 for at least 5 consecutive milliseconds), and cumulative distributions were compared using two-sample Kolmogorov-Smirnov tests with Bonferroni correction (α = 0.05/3 = 0.0167 per cluster).

**Figure 6.**
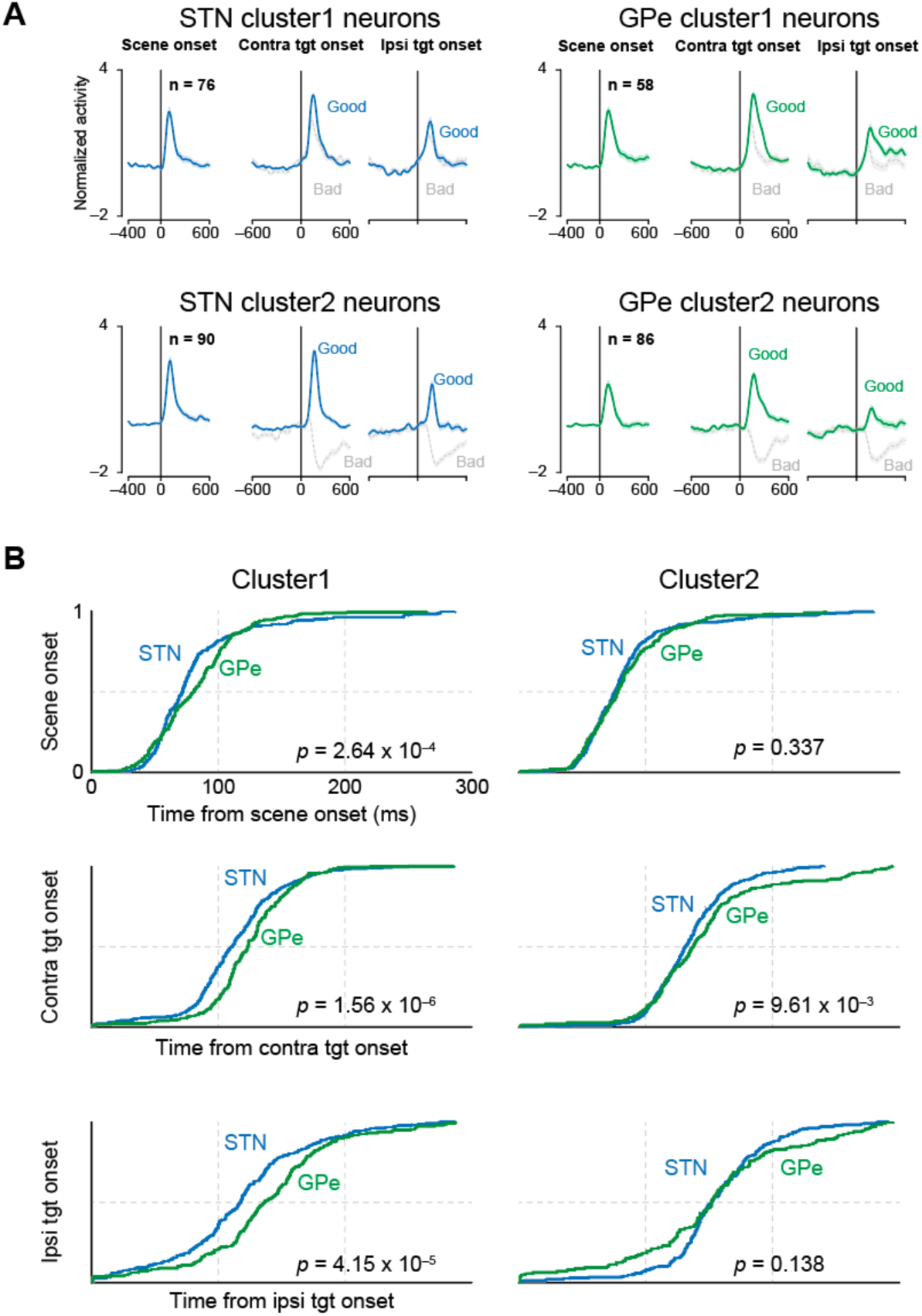
Comparison of response latencies between STN and GPe neurons. (A) Population-averaged normalized activity (z-scored) for STN clusters (left column; present study) and functionally analogous GPe clusters (right column) recorded during the same task paradigm.^31^ Colored traces indicate good-object conditions (blue: STN; green: GPe), which were used for the latency comparison in (B). Gray traces indicate bad-object conditions. Activity is shown aligned to scene onset (left), contralateral target onset (center), and ipsilateral target onset (right). The number of neurons in each population is indicated (n). (B) Cumulative distribution functions (CDFs) of neuronal response latency for STN (blue) and GPe (green) populations. Response latency was defined as the first time point (100–300 ms post-event) at which z-scored activity exceeded 2.0 for at least 5 consecutive milliseconds (see Methods). Populations were matched based on analogous response profiles: STN Cluster 1 with GPe Cluster 1 (left column), and STN Cluster 2 with GPe Cluster 2 (right column). *P*-values are from two-sample Kolmogorov-Smirnov tests (Bonferroni-corrected significance threshold within each cluster: α = 0.05/3 = 0.0167). STN response latencies were significantly shorter than GPe latencies for all three conditions in Cluster 1, and for contralateral good targets in Cluster 2.

For Cluster 1, STN response latencies were significantly shorter than GPe latencies across all three conditions: scene onset (*P* = 2.64 × 10⁻⁴), contralateral good target onset (*P* = 1.56 × 10⁻⁶), and ipsilateral good target onset (*P* = 4.15 × 10⁻⁵; Figure 6B, left). For Cluster 2, STN preceded GPe for contralateral good target onset (*P* = 9.61 × 10⁻³), but not for scene onset (*P* = 0.337) or ipsilateral good target onset (*P* = 0.138; Figure 6B, right). The condition-selective temporal precedence in Cluster 2 is consistent with its functional profile, as this cluster shows strong value-dependent spatial selectivity specifically for contralateral good objects, and it is in this preferred condition that the temporal precedence is most evident.

These results demonstrate that STN responses systematically preceded those of functionally matched GPe populations, particularly under conditions that engage the preferred stimulus of each cluster. Combined with the close correspondence in response profiles (Figure 6A) and the dense anatomical connectivity between the ventral STN and dorsal GPe,^32^ these findings are consistent with a feedforward excitatory influence from the STN to the GPe during value-based action selection.

### Single-Neuron and Population-Level Encoding of Choice

To quantify how strongly individual STN neurons encoded the monkey’s choice over time, we performed a trial-by-trial analysis using a sliding-window general linear model (GLM) for contralateral trials (Figure 7A). In this analysis, the mean neuronal activity within a 50-ms window was regressed against the upcoming choice (’accept’ = 1, ‘reject’ = –1). The resulting heatmaps of β coefficients in Figure 7A revealed distinct patterns across clusters. A large majority of neurons in Cluster 1, and nearly all in Cluster 2, showed positive β coefficients, robustly encoding the ‘accept’ choice. In contrast, most neurons in Cluster 3 exhibited negative β coefficients, consistent with encoding the ‘reject’ choice.

**Figure 7.**
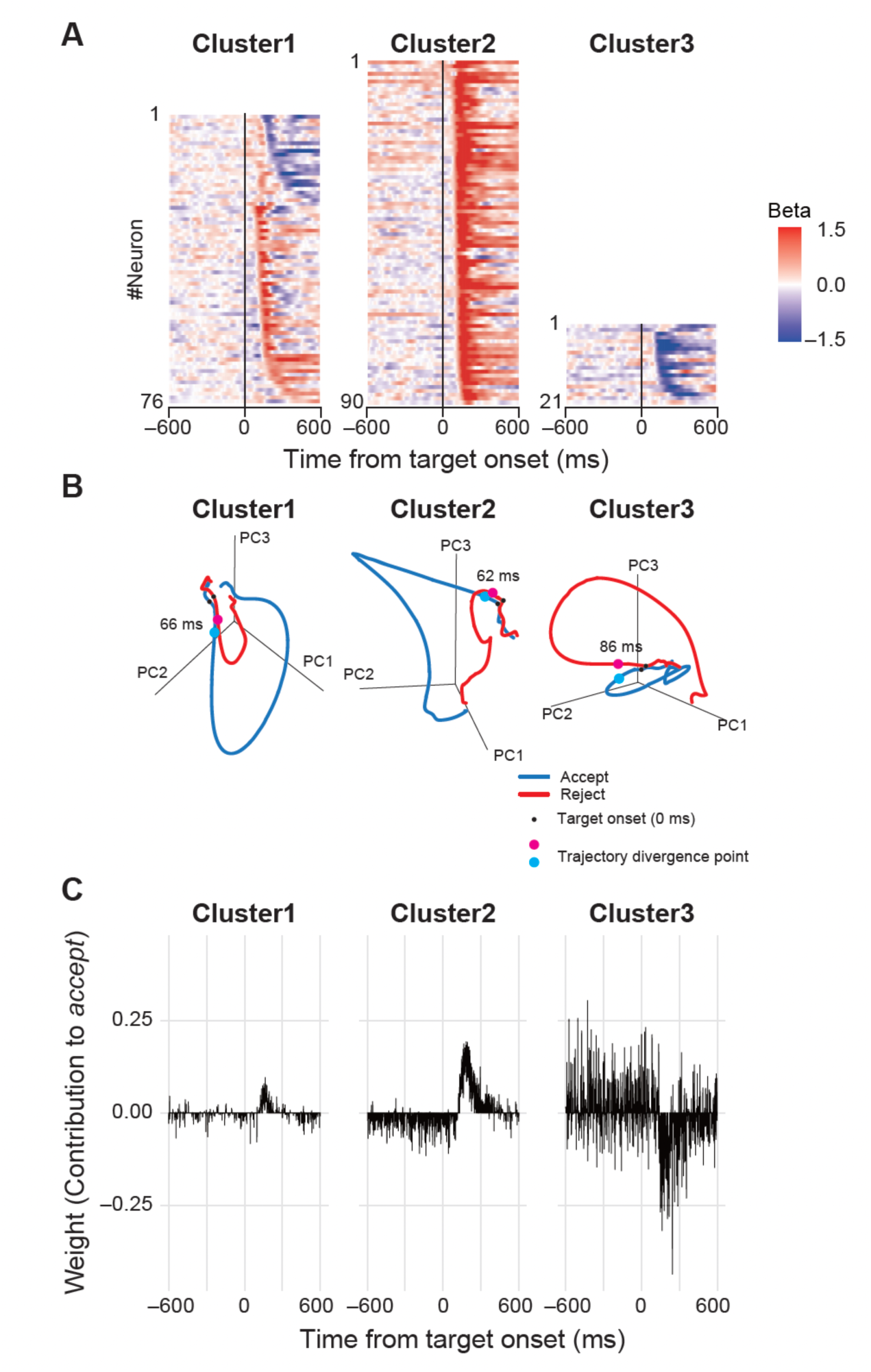
Single-neuron and population-level encoding of choice. (A) Single-neuron choice encoding. Heatmaps show the time course of β coefficients from a sliding-window GLM for each neuron. Red indicates a positive β (encoding ‘accept’), while blue indicates a negative β (encoding ‘reject’). Neurons within each cluster are sorted by their response preference and latency. (B) Population activity trajectories. The evolution of the population activity for each cluster is visualized as a trajectory in a 3D state space derived from PCA. Traces for ‘accept’ (blue) and ‘reject’ (red) choices are shown. The black dot indicates target onset (t = 0), and the cyan and magenta dots indicate the time at which the two trajectories became statistically separable. (C) Population decoding weights for each cluster. The time course of weights from three separate Elastic Net classifiers, each trained to predict choice using the population activity from neurons within Cluster 1, 2, and 3, respectively. A positive weight indicates that neuronal activity at a given time point contributes to predicting an ‘accept’ choice, whereas a negative weight contributes to predicting a ‘reject’ choice.

To understand how these individual signals combined and evolved collectively, we next visualized the population activity using PCA on the trial-averaged spike density functions. This approach visualizes the population activity as neuronal trajectories in a low-dimensional state space (Figure 7B).

In all clusters, the trajectories for ‘accept’ and ‘reject’ choices began to separate into distinct paths shortly after the target appeared. We quantified this separation and found that the neuronal representations diverged remarkably early (62–86 ms post-target onset), well before the average saccadic reaction times. This indicates that the STN population activity reflects the outcome of the decision-making process itself, rather than simply encoding the subsequent motor command.

To quantify the predictive power of the STN population, we performed a decoding analysis using an Elastic Net classifier (Figure 7C). To obtain an unbiased performance estimate and prevent data leakage from non-independent trials, we employed a nested, 10-fold cross-validation scheme where folds were grouped by neuron (see Methods). When analyzing the pooled data from both monkeys, the model predicted the action choice with an *AUC* = 0.747 and, to a lesser extent, also predicted RTs (cross-validated *R²* = 0.062).

To assess the relative importance of each cluster, we performed both ablation and single-cluster analyses. The results revealed a critical and dominant role for Cluster 2. In the ablation analysis, removing Cluster 2 from the model caused performance to drop precipitously to an *AUC* of 0.558, a level barely above chance. Conversely, removing either Cluster 1 (*AUC* = 0.807) or Cluster 3 (*AUC* = 0.806) slightly improved the overall performance. This conclusion was strengthened by the single-cluster analysis. A model using only Cluster 2 neurons achieved a remarkably high performance of *AUC* = 0.918, while models using only Cluster 1 (*AUC* = 0.614) or Cluster 3 (*AUC* = 0.607) neurons performed substantially worse.

Crucially, these key findings were highly consistent when the analysis was repeated for each monkey individually. For both Monkey C and Monkey S, a model using only Cluster 2 neurons achieved high accuracy (*AUC* = 0.885 and 0.916, respectively), while removing Cluster 2 consistently caused a dramatic drop in performance (*AUC* = 0.519 and 0.573, respectively).

Taken together, these population-level analyses provide strong quantitative support for a functional division of labor within the STN. The predictive information for value-based choice is not distributed evenly across populations but is instead overwhelmingly concentrated in the Cluster 2 neurons, which powerfully signal the facilitation of a desired action.

## Discussion

In the present study, we investigated the role of the primate STN in value-based decision-making. Our findings reveal that the STN is not a functionally homogeneous structure but is composed of distinct functional groupings. Critically, the vast majority of task-related neurons exhibited activity consistent with stimulus evaluation and action facilitation, challenging the traditional view of the STN as primarily a brake on motor output. These results suggest the STN plays a multifaceted role, with a predominant function in the facilitation of desired actions toward valued targets, while also containing a smaller component that signals the negative value of unrewarded stimuli.

### Functional Organization of STN Neurons during Value-Based Decisions

The present results reveal a functional organization within the ventral STN during value-based decision-making. The two facilitative populations (Clusters 1 and 2) constituted the majority of task-related neurons (approximately 89%), consistent with a long history of studies reporting movement-related increases in STN activity across species and task contexts.^11–17^ Rather than characterizing the STN as a simple “accelerator” or “brake,” however, our findings demonstrate that these facilitative neurons are functionally heterogeneous and operate predominantly in a stimulus-driven evaluative mode. The temporal alignment analysis showed that the majority of facilitative neurons were target-locked (Cluster 1: 68%, Cluster 2: 76%), indicating that their activity reflects the evaluation of visual stimuli rather than direct motor commands. This interpretation is reinforced by the spatial selectivity of these neurons, specifically their contralateral preference for valued targets, which mirrors the properties of visual rather than motor responses.^24^ The saccade-locked minority within these clusters represents a subpopulation whose activity is more closely related to movement execution, suggesting that the STN covers a functional range from stimulus-driven evaluation to motor-related processing.

The population-level relationship between neuronal onset latency and saccadic reaction time was statistically significant but modest in magnitude (β = 0.036 and 0.057). The temporal alignment analysis provides a more precise interpretation of this result. The target-locked majority of neurons contributes little to the onset-RT association because their timing is linked to the sensory event, while the saccade-locked minority of neurons drives the significant relationship because their timing varies together with movement execution. The strong correlation between Locking Index and RT-onset slope (ρ = 0.631) confirms that these two measures, one based on variability and the other on regression, converge on a consistent characterization of where each neuron falls along the sensory-motor range. Furthermore, our population-level choice analysis showed that the neuronal trajectories for ‘accept’ and ‘reject’ choices diverged well before movement onset, indicating that the STN is involved in the decision-making process itself, rather than simply relaying a pre-determined motor command. These properties, including predominantly target-locked responses with value and spatial modulation, are consistent with the facilitative function reported in rodents, where ‘type 1’ STN neurons were essential for sustaining locomotion,^15^ suggesting that the facilitative role of the STN is a conserved principle that extends to value-based decision-making in primates.

The third population (Cluster 3) was initially characterized as potentially consistent with the classical suppressive role of the STN. However, two converging lines of evidence argue against a motor suppression interpretation. First, 95% of Cluster 3 neurons were target-locked, indicating that their activity was linked to the sensory event rather than to the motor response. Second, Cluster 3 activity did not differ between return (saccade execution) and stay (fixation maintenance) rejection strategies for bad objects, demonstrating that the signal is independent of the motor strategy used to avoid the unrewarded target (Figure S6). Together, these findings indicate that Cluster 3 encodes a stimulus-driven negative value signal rather than a motor brake.

This distinction is important because the classical braking function of the STN has been proposed primarily based on stop-signal tasks, in which subjects must reactively cancel an already-initiated movement. Our value-based choice task is fundamentally different, as monkeys evaluate visual objects and select an appropriate action based on learned value. The most common response to bad objects was a return saccade, which involves the execution, not the suppression, of a movement. The finding that Cluster 3 activity did not differ between return and stay strategies further dissociates this signal from motor cancellation. Rather than a reactive brake triggered by a stop signal, Cluster 3 appears to encode a proactive evaluation of stimulus value that is computed independently of the subsequent motor plan.

To further validate the functional architecture identified by our clustering analysis, we also applied PCA. This analysis revealed a consistent organizational principle, namely a primary division of the STN population into two major facilitative profiles. A purely negative value-coding group analogous to Cluster 3 was not isolated as a separate component, suggesting that while this population is functionally distinct, it may not define a separable class of neurons at the population level. Rather than operating as a simple “brake” or “accelerator,” the STN appears to function as a value-based action selector, with the majority of neurons evaluating stimuli to facilitate the selection of rewarded actions and a smaller population signaling the negative value of unrewarded stimuli.

### Functional topography and limitations of spatial sampling

Our recordings were concentrated in the ventral portion of the STN, as confirmed by one-sample t-tests comparing recording coordinates against the STN centroid in both monkeys (Figure S3B). This region overlaps with the oculomotor and associative territories defined by anatomical tracing studies^25,26^ and is consistent with the saccade-based nature of our behavioral task. Within this ventral territory, the three functional clusters showed no significant topographic segregation (Kruskal-Wallis tests, all FDR-corrected *P* > 0.06), indicating that neurons with distinct functional properties are spatially intermingled. Spike waveform analysis further revealed a significant difference in peak-to-peak duration between Cluster 1 and Cluster 3 (*P* = 0.010), which may reflect differences in physiological properties across clusters. However, because extracellular waveform measures can be influenced by recording geometry and signal quality, this finding should be considered supportive but not definitive evidence for distinct neuronal subtypes.

An important limitation of the present study is that our conclusions are specific to the ventral STN and may not generalize to the dorsal sensorimotor territory. The dorsal STN receives inputs from motor cortex and is associated with limb and trunk movements.^25^ Studies recording from the dorsal STN during limb movement tasks have reported a different balance of movement-related responses,^36,37^ and it remains possible that the proportion of facilitative versus negative value-coding neurons differs across STN subregions. Future studies combining recordings across the full dorsoventral extent of the STN with tasks that engage both oculomotor and skeletomotor systems will be needed to determine whether the functional organization described here is specific to the ventral territory or reflects a general principle of STN processing.

### Comparison with previous classifications of STN activity

Matsumura et al. (1992) classified STN neurons recorded during visually guided saccades into visual and saccade-related categories based on their temporal relationship to the stimulus and movement.^24^ Our temporal alignment analysis provides a quantitative framework for this classification. The target-locked majority in Clusters 1 and 2 (68% and 76%, respectively) corresponds to their visual responses, while the saccade-locked minority corresponds to their saccade-related activity. The spatial selectivity of our facilitative clusters, with contralateral preferences for valued targets, is also consistent with the directional tuning reported in that study.

However, our results extend this earlier framework in two important ways. First, our value-based task reveals that the visual responses described in their classification are not simply sensory signals but carry information about learned object value. The contralateral preference in Cluster 1 was present for both good and bad objects, while in Cluster 2 it was restricted to good objects, indicating that value modulates spatial coding in a cluster-specific manner.

Second, Cluster 3 represents a functional category not identified in the earlier study. This population is almost exclusively target-locked (95%) and non-directional, encoding a negative value signal upon presentation of unrewarded objects. Because their task did not manipulate stimulus value, this category would not have been distinguishable from other visually responsive neurons in their paradigm. Together, these findings suggest that the functional organization of the STN during value-based decisions can be described along three dimensions: temporal alignment (target-locked vs saccade-locked), spatial selectivity (contralateral vs non-directional), and value coding (good-preferring, bidirectional, or bad-preferring).

### Circuit mechanisms underlying STN-mediated action selection

How does excitatory STN activity facilitate movement? Our data, combined with recordings from the GPe^31^ and SNr^34^ obtained during the identical paradigm in the same monkeys, suggest a circuit model in which different STN populations access distinct output pathways.

We propose that the facilitative clusters (Clusters 1 and 2) promote action primarily through the STN–GPe–SNr disinhibition pathway. In this model, STN excitation could increase activity in GPe neurons, which in turn inhibit SNr neurons, releasing downstream motor structures such as the superior colliculus and thalamus from tonic inhibition. This proposal is supported by the close correspondence between the response profiles of our facilitative STN clusters and functionally analogous GPe populations (Figure 6A), and by the systematic temporal precedence of STN responses over GPe responses (Figure 6B).

Furthermore, the ventral STN and dorsal GPe, where our recordings were concentrated, are densely interconnected in primates.^32^ The response properties of the facilitative GPe populations resemble those of prototypical GPe neurons, consistent with findings in rodents showing that STN projections preferentially target prototypical neurons implicated in movement promotion.^30^

The activity pattern observed in the lateral SNr provides an additional constraint on this model. In our SNr recordings,^34^ the vast majority of lateral SNr neurons showed decreased activity for good objects and increased activity for bad objects. Notably, almost no SNr neurons showed increased activity for good objects. If the facilitative STN clusters projected directly to the SNr, their excitation for good objects would be expected to increase SNr activity, which was not observed. This absence is consistent with the facilitative clusters influencing the SNr predominantly indirectly via the GPe. Specifically, STN excitation could increase GPe activity, thereby enhancing GPe–SNr inhibition, producing the observed decrease in SNr activity for good objects and thereby facilitating saccades to valued targets. In contrast, the increased activity of Cluster 3 for bad objects, transmitted via the direct excitatory STN–SNr pathway, would produce the observed increase in SNr activity for bad objects, which in turn suppresses downstream motor output. Although the downstream consequence of this pathway is motor suppression, the signal encoded by Cluster 3 itself is stimulus-driven and reflects object value rather than a motor cancellation command.

Under this model, the indirect inhibitory signal from the facilitative clusters (via the GPe) and the direct excitatory signal from Cluster 3 converge at the SNr, where they are integrated to produce the final motor output. This convergence accounts for the bidirectional value coding observed in the lateral SNr and provides a unified circuit framework for how the STN contributes to value-based action selection. It is important to acknowledge that the STN is not the sole source of excitatory input to the GPe; a direct corticopallidal pathway also provides significant excitatory drive,^38–40^ and our findings do not exclude its potential contribution to the facilitative GPe activity. Furthermore, these pathway assignments remain hypothetical, as the present data do not identify the projection targets of individual neurons. Techniques such as antidromic stimulation or projection-specific recordings will be needed to test this model directly.

### Clinical implications

The recognition of distinct facilitative and negative value-coding pathways through the STN has clinical implications. The complex effects of STN deep brain stimulation (DBS) in Parkinson’s disease, which can improve motor facilitation as well as inhibition,^41–43^ are difficult to explain using conventional models that consider only the STN–SNr suppressive pathway. Our findings suggest that the therapeutic effects of DBS may be mediated through both the facilitative STN–GPe–SNr pathway and the direct STN–SNr pathway. DBS applied to the STN would affect both pathways simultaneously, potentially explaining why stimulation can produce improvements in both movement initiation and action suppression. Incorporating this dual-pathway model into existing theory may be important for a more complete understanding of both normal basal ganglia function and the pathophysiology of related movement disorders.

### Limitations and Future Directions

The present study has several limitations. First, our findings are correlational; while we demonstrate a strong link between STN activity and behavior, we have not established a causal relationship. Second, the functional analogy between STN and GPe populations does not constitute direct proof of a monosynaptic circuit. It remains possible that other excitatory inputs, for instance from the cortex, also contribute to the facilitative responses observed in the GPe.

Future studies employing circuit-dissection tools are essential to address these limitations. Establishing causality will require techniques such as optogenetics or chemogenetics (e.g., DREADDs) to selectively manipulate the activity of specific STN clusters (e.g., the facilitative populations) while observing the impact on both GPe neuronal activity and behavior. Furthermore, pathway-specific viral tracing methods will be necessary to anatomically map the projections from each functional STN cluster to their precise targets within the GPe and other basal ganglia nuclei. Such experiments will be crucial not only for validating the proposed STN-GPe facilitative circuit but also for delineating its functional role relative to the canonical direct pathway, which also contributes to action promotion. Ultimately, integrating these novel pathways into established models of the basal ganglia is an important and necessary step toward a more complete understanding of how the brain controls voluntary movement.

## Acknowledgments

This research was supported by the Intramural Research Program of the National Institutes of Health, National Eye Institute (ZIA EY000415 (O.H.) and ZIA EY000511 (R.J.K)). The contributions of the NIH authors were made as part of their official duties as NIH federal employees, are in compliance with agency policy requirements, and are considered Works of the United States Government. However, the findings and conclusions presented in this paper are those of the authors and do not necessarily reflect the views of the NIH or the U.S. Department of Health and Human Services. MRI scanning was conducted at the Neurophysiology Imaging Facility Core (National Institute of Mental Health, National Institute of Neurological Disorders and Stroke, and National Eye Institute). We thank D. Parker, H. Warnock, G. Tansey, K. Allen-Worthington, A.M. Nichols, D. Yochelson, J. Fuller-Deets, and M. Robinson for their technical assistance.

## Author contributions

Conceptualization, A.Y., O.H.; Methodology, A.Y., R.J.K; Software, A.Y.; Validation, A.Y.; Formal Analysis, A.Y.; Investigation, A.Y.; Resources, O.H.; Data Curation, A.Y.; Writing – Original Draft, A.Y.; Writing – Review & Editing, A.Y., R.J.K; Visualization, A.Y.; Supervision, O.H.; Funding Acquisition, O.H., R.J.K

## Methods

### Subjects

All experimental procedures were approved by the National Eye Institute Animal Care and Use Committee and conducted in accordance with the Public Health Service Policy on Laboratory Animal Care. Two male rhesus macaques (Macaca mulatta, 8-10 kg), designated as Monkeys C and S, served as subjects. These animals were also used in our previous studies.^31,33,34^ Surgical procedures were performed under isoflurane anesthesia and aseptic conditions to implant a plastic head holder and recording chambers. After a recovery period, the monkeys were trained to perform oculomotor tasks. During experimental sessions, the monkeys’ heads were held in a fixed position, and we monitored their eye movements at 1000 Hz using an infrared eye-tracking system (EyeLink 1000, SR Research). To maintain motivation, fluid intake was regulated throughout the experimental period. Detailed descriptions of the surgical and postoperative care have been published previously.^33^

### Behavioral Task and Visual Stimuli

Experiments took place in a light- and sound-attenuated room, with visual stimuli presented on a screen via an LCD projector (PJ658, ViewSonic). Task control and data acquisition were managed by custom C++ software. The choice task paradigm was identical to that used in our previous work.^33^

In each session, one of six scene-object sets was chosen at random. Each set contained four scenes, and each scene was associated with a “good” object linked to a liquid reward and a “bad” object yielding no reward. For Scenes 1 and 2, object-reward contingencies were stable, while for Scenes 3 and 4, the values were reversed. This flexible-value design allowed us to dissociate neuronal responses to object value from those driven by low-level visual features.

A typical trial began with a 1000-ms scene presentation, followed by a 700-ms central fixation period. Subsequently, a good or bad object appeared at one of six peripheral locations (15° eccentricity). To accept the object, the monkey had to make a saccade to it and maintain fixation for at least 400 ms, which resulted in a reward (0.4 mL of juice) for a good object. To reject an object, the monkey could use one of three strategies: a brief saccade to the object followed by a return to center (“return”), maintaining central fixation (“stay”), or a saccade to a location away from both the object and the center (“other”).

### MRI and Localization

Following chamber implantation, MRI scans were performed to map anatomical structures relative to the recording grid. All MR images were acquired under anesthesia to avoid head motion artefact. The monkey’s head was fixed in an MRI-compatible stereotaxic frame. Atropine (0.05 mg/kg, i.m.) was initially injected and ketamine (10 mg/kg, i.m.) and dexmedetomidine (0.01 mg/kg, i.m.) were used for induction of anesthesia. Additional ketamine (5 mg/kg, i.m.) and dexmedetomidine (0.01 mg/kg, i.m.) were injected for maintenance. Recording sites were localized using a 3-T MRI system (MAGNETOM Prisma, Siemens) to acquire three-dimensional T1-weighted (T1w) and T2-weighted (T2w) sequences at 0.5-mm isotropic resolution.

To enhance the visualization of the STN, we employed quantitative susceptibility mapping (QSM), which provides superior contrast for iron-rich subcortical structures compared to conventional imaging.^44–46^ QSM images were reconstructed from phase images acquired with a 3D multi-echo gradient echo sequence (repetition time: 50 ms; echo times: 3.7, 10.1, 16.7, 23.4, 30.0, 36.6, 43.2 ms).^46^ The reconstruction pipeline, implemented in MATLAB 2019 using the morphology-enabled dipole inversion toolbox, included phase unwrapping, background field removal, and dipole inversion to generate the final susceptibility maps.^47^

To precisely locate the grid holes, a high-resolution T1w MPRAGE sequence (0.33 × 0.33 × 0.35 mm³ voxel size) was acquired while the grid was filled with a gadolinium-based contrast agent. For recording guidance, we created fused images combining this high-resolution scan with the QSM or conventional T1w images. To illustrate the recording locations (Figure S3B), the STN was automatically segmented on each monkey’s T1w image using an AFNI-based pipeline, which aligned the native brain to a standard macaque template (NMT v2.0) and then inversely transformed the Subcortical Atlas of the Rhesus Macaque (SARM).^48–51^ The accuracy of this segmentation was confirmed by comparison with the QSM images, which provide superior visualization of the STN structure compared to standard T1w and T2w images.^46,52^ Final 3D visualizations were created using 3D Slicer (v5.2.2)^53^, Blender (v3.5), and the matplotlib Python library (v3.10.0).^54^

### Electrophysiology

We began single-unit recordings in the STN after the monkeys achieved stable task performance, defined as >90% accuracy in discriminating between good and bad objects. We used tungsten microelectrodes (1–9 MΩ; FHC; Alpha Omega) that were advanced through stainless steel guide tubes using a hydraulic micromanipulator (Narishige). The recorded neuronal signals were amplified, bandpass filtered (0.3–10 kHz; A-M Systems), and digitized at 40 kHz. A neuron was selected for recording if it exhibited clear task-related modulation following target onset and could be well-isolated from background noise and other units. Single neurons were isolated online using a voltage-time window discriminator, where units were separated based on both their waveform amplitude and specific shape using a combination of inclusion and exclusion windows.

## QUANTIFICATION AND STATISTICAL ANALYSIS

All data were preprocessed and analyzed using MATLAB 2022b and R (R Core Team).^55^ The sample size was determined based on previous studies recording from the primate STN.^24^

### Behavioral Data Analysis

Saccade onsets were detected when eye velocity surpassed 40°/s. Reaction times (Figure S1A) were calculated for initial saccades to objects and were compared between conditions using the Welch t-test. The proportion of “stay” responses between stable- and flexible-value scenes was compared using Fisher’s exact test.

### Neuronal Data Analysis

Spike trains were aligned to scene, target, and saccade onsets. Peristimulus time histograms (PSTHs) were constructed with 1-ms bins and smoothed with a Gaussian kernel (σ = 20 ms). For population-level analyses, neuronal activity was z-transformed by subtracting the mean firing rate during a 500-ms pre-event baseline period and dividing by the standard deviation of that baseline activity.^56,57^

### Clustering Analysis

Neurons were classified using k-means clustering based on their average z-scored firing rates in a 200-ms window (100–300 ms) after the presentation of contralateral good and bad objects. To determine the optimal number of clusters (K), we used the silhouette method. We simulated silhouette values 5,000 times for K ranging from 2 to 6 and selected the K that yielded the highest average value (Figure S2A).

### Principal Component and Clustering Analysis

To validate the functional groupings identified by the window-based method and to characterize the neural population in a more data-driven manner, we performed PCA on the trial-averaged neural activity.

The input matrix for the PCA was constructed from the Z-scored activity on contralateral trials. To treat the responses to different stimulus values independently, the average responses to ‘good’ and ‘bad’ objects for each neuron were stacked as separate entries, resulting in a matrix with 374 rows (187 neurons × 2 conditions). This analysis was performed on a short window of activity (0 to 300 ms post-stimulus). PCA was performed using the *prcomp* function in R, with the data for each time point centered and scaled. A cumulative variance plot was generated to determine the number of components to retain for subsequent analysis, which indicated that the first three PCs collectively captured the majority (>60%) of the structured variance in the data (Figure S4B).

To subsequently group neurons based on the functional properties revealed by these top three PCs, we applied a k-means clustering algorithm. The feature space for clustering was derived from the PCA scores (weights). For each of the 187 neurons, we extracted its scores on the top three principal components (PC1-3) for both the ‘good’ and ‘bad’ conditions. These were combined to form a six-dimensional feature vector for each neuron. K-means clustering (*kmeans* function in R) was then applied to this 187-neuron × 6-feature matrix.

### Linear Mixed-Effects Models (LMMs)

To compare normalized neuronal activity across different conditions (Figures 3 and 4), we used LMMs to reduce type I errors and better represent the data structure.^58^ These models included fixed effects for the experimental variables and their interactions, with random intercepts for monkey and for neuron nested within monkey to account for non-independence of the data. The general structure of the most complex model, using R’s formula notation, was:

NormalizedActivity ∼ Scene ∗ Value ∗ Direction + (1|MonkeyID) + (1|MonkeyID:NeuronID)

Model significance was assessed using a parametric bootstrap method (10,000 iterations) comparing the full model to a null model containing only the random effects. Significant main effects or interactions were followed by post-hoc pairwise t-tests with Bonferroni correction. These analyses were performed in RStudio using the lme4, pbkrtest, and emmeans packages.^59–62^

### Temporal alignment analysis

To determine whether the response of each neuron was time-locked to the sensory event (target onset) or to the motor response (saccade onset), we adapted the variance-based framework of the previous study.^35^ For each neuron, we identified the single-trial response onset latency as the first time point within a 100–300 ms post-target window at which the z-scored firing rate exceeded a threshold of 2.0 for at least 5 consecutive milliseconds. Because onset detection was intentionally constrained to this early post-target window (100–300 ms), the Locking Index primarily reflects the temporal alignment of the early task-evoked response component and is not intended to capture later movement-related activity. Neurons for which onset could not be detected on a sufficient number of trials were excluded from this analysis.

For each neuron, we computed the standard deviation of single-trial onset latencies measured relative to target onset (SD_target) and relative to saccade onset (SD_saccade). A Locking Index (LI) was then defined as:

LI = (SD_target − SD_saccade) / (SD_target + SD_saccade)

This index ranges from −1 (purely target-locked) to +1 (purely saccade-locked), with LI = 0 indicating equal variability relative to both events. Neurons with LI < 0 were classified as target-locked; neurons with LI > 0 were classified as saccade-locked (Figure 5A, B).

To validate this classification with an independent metric, we performed a per-neuron linear regression of saccadic reaction time on response onset latency (measured relative to target onset). The resulting slope quantifies how strongly the onset timing of each neuron covaries with movement initiation: a slope near 0 is expected for target-locked neurons whose onset timing is independent of RT, while a slope approaching 1 is expected for saccade-locked neurons whose onset timing shifts with saccade initiation. The convergence between the Locking Index and the RT-onset slope was assessed using Spearman rank correlation, computed across all neurons and within each cluster separately (Figure 5C).

### Response latency comparison across STN clusters

To compare the speed of neuronal responses across the three functional clusters, we measured single-neuron response latencies for three task events, scene onset, contralateral target onset, and ipsilateral target onset. Response latency was defined as the first time point within a 100–300 ms post-event window at which z-scored activity exceeded a threshold of 2.0 for at least 5 consecutive milliseconds. For scene onset, latency was measured identically across all clusters. For target onset conditions, latency was measured using the preferred stimulus of each cluster to ensure reliable onset detection. Good objects were used for Clusters 1 and 2, and bad objects were used for Cluster 3. Neurons for which onset could not be detected were excluded from the comparison for that condition. Data were pooled across scenes 1–4, as individual neurons showed consistent response patterns across scenes. The cumulative distributions of response latencies were compared between all cluster pairs using two-sample Kolmogorov-Smirnov tests with Bonferroni correction for three pairwise comparisons (corrected significance threshold, α = 0.05/3 = 0.0167).

### Comparison of response latencies between STN and GPe populations

To test the hypothesis that STN activity exerts a feedforward excitatory influence on the GPe, we compared neuronal response latencies between STN populations recorded in the present study and functionally analogous GPe populations from our previously published study using the same task paradigm (Yoshida and Hikosaka, 2025).^31^ The GPe recordings were obtained from the same two monkeys used in the present study (Monkey Cr and Monkey Sp in previous study,^31^ corresponding to Monkey C and Monkey S in the present study, respectively). Both datasets were collected while the monkeys performed the same value-based sequential choice task with the same visual stimuli and reward contingencies, allowing direct comparison of neuronal response latencies between the two structures.

STN and GPe populations were matched based on their analogous response profiles. STN Cluster 1 was paired with GPe Cluster 1 (both showing excitatory responses to good and bad objects with a good-object preference), and STN Cluster 2 was paired with GPe Cluster 2 (both showing bidirectional value coding with excitation for good objects and suppression for bad objects). For each paired comparison, response latencies were measured for good-object conditions using the same criteria as described above (z-score threshold of 2.0 for at least 5 consecutive milliseconds within a 100–300 ms post-event window). Cumulative distributions were compared using two-sample Kolmogorov-Smirnov tests for three conditions (scene onset, contralateral good target onset, and ipsilateral good target onset), with Bonferroni correction applied within each cluster pair (corrected significance threshold: α = 0.05/3 = 0.0167).

### Relationship between Neuronal Activity and Reaction Time

To examine the trial-by-trial relationship between neuronal activity and RT, we focused on ‘accept’ trials for contralateral good objects. For each trial, we calculated the z-scored SDF (Spike Density Functions) and extracted three parameters: 1) Response Onset Latency (the first time point after target onset where the z-score exceeded 2), 2) Peak Response Magnitude (the maximum z-score within a 150-ms window following the response onset), and 3) Response Slope (the rate of change in z-score from onset to peak).

To statistically assess the relationship between each neuronal parameter and RT, we fitted separate LMMs. These models included the neuronal parameter as a fixed effect to predict RT, with random intercepts for MonkeyID and NeuronID to account for non-independence of the data. The general structure of the models, consistent with our R script implementation, was:

RT ∼ NeuronalParameter + (1 | MonkeyID) + (1 | MonkeyID:NeuronID)

The significance of the fixed effect was assessed based on the *P*-value derived from Satterthwaite’s method for approximating degrees of freedom. A Bonferroni correction was applied to account for the nine separate models fitted, resulting in a significance threshold of *P* < 0.0056. These analyses were performed in RStudio using the *lme4* and *lmerTest* packages. For visualization in Figure S5, each scatter plot was overlaid with the predicted marginal effect line and its 95% confidence interval derived from the LMM. The unstandardized regression coefficient (β) and its *P*-value from the model were displayed on each plot.

### Sliding-window General Linear Model (GLM)

To quantify how the choice signal of individual neurons evolved over time, we performed a trial-by-trial analysis using a sliding-window GLM on contralateral trials.^59^ For each neuron, a 50-ms analysis window was moved in 1-ms steps across the trial epoch. At each step, we fitted a linear model (*lm* function in R) of the form:

MeanActivity = β_0_ + β_Action_ * Action + ɛ

where Action was coded as 1 for ‘accept’ and −1 for ‘reject’. The β coefficient for the Action term was extracted at each time point. For visualization (Figure 7A), these β coefficients were smoothed (21-ms moving average) and sorted by response preference and latency.

### Population-level Analysis using PCA

To visualize population dynamics, we used PCA for dimensionality reduction. For each neuron, spike density functions (SDFs; σ = 20 ms) from contralateral trials were baseline-corrected. Trial-averaged SDFs for ‘accept’ and ‘reject’ conditions were combined to fit a single PCA model. The population activity for each condition was then projected onto the first three PCs, which captured the largest amount of variance in the data. To statistically determine the time of choice signal emergence, we calculated the Euclidean distance between the two trajectories in PC space at each time step. The divergence time was defined as the first time point post-target where this distance exceeded three standard deviations above the mean baseline distance.

### Population Decoding using Elastic Net

To quantify the predictive power of the STN population, we used an Elastic Net regularized regression approach. For each trial, spike counts in 1-ms bins (–600 to 600 ms from target onset) across all neurons were concatenated into a feature vector.^60^ All feature standardization (z-scoring) was fit exclusively on the training data within each cross-validation fold and then applied to the corresponding test sets to prevent information leakage.

To obtain an unbiased estimate of model performance and prevent data leakage from non-independent trials, we assessed performance using a nested, 10-fold cross-validation procedure, grouped by neuron. In the outer loop, the dataset was split into 10 folds such that all trials from a given neuron belonged to the same fold. For each iteration of the outer loop, 9 folds were used for training, and the remaining fold was held out for testing. Within the training set, an inner 10-fold cross-validation (also grouped by neuron) was performed to select the optimal regularization parameter (*λ*) that maximized the area under the ROC curve (*AUC*) for choice prediction or minimized mean-squared error (MSE) for RT prediction. A final model was then trained on the entire outer training set using this optimal λ and evaluated on the held-out test fold. The overall performance metric (*AUC* or *R²*) was calculated from the aggregated predictions from all 10 outer test folds. We set the Elastic Net mixing parameter α to 0.5 a priori, while λ was selected in the inner cross-validation loop.

To assess the importance of each cluster, we performed an ablation analysis (systematically removing one cluster at a time) and tested the performance of each cluster individually, applying the same nested, grouped cross-validation procedure to each subset of data. The statistical significance of the single-cluster *AUCs* was determined via a permutation test (2,000 iterations). In each iteration, choice labels were shuffled, and the entire nested, grouped cross-validation procedure was repeated to generate an accurate null distribution of *AUCs*. This entire set of decoding analyses was conducted first on the data pooled from both monkeys and then repeated on the data from each monkey individually to confirm the consistency of the findings.

## Data and Code Availability

The datasets generated and analyzed during the current study have been deposited at Zenodo and are publicly available as of the date of publication at https://doi.org/10.5281/zenodo.17860356. The DOI is also listed in the Key Resources Table.

Any additional information required to reanalyze the data reported in this paper is available from the lead contact upon request.

## Declaration of generative AI and AI-assisted technologies in the writing process

During the preparation of this work the authors used Gemini 3.1 in order to check typos and grammatical errors and improve readability. After using this tool, the authors reviewed and edited the content as needed and take full responsibility for the content of the published article.

---

**Table S1.**
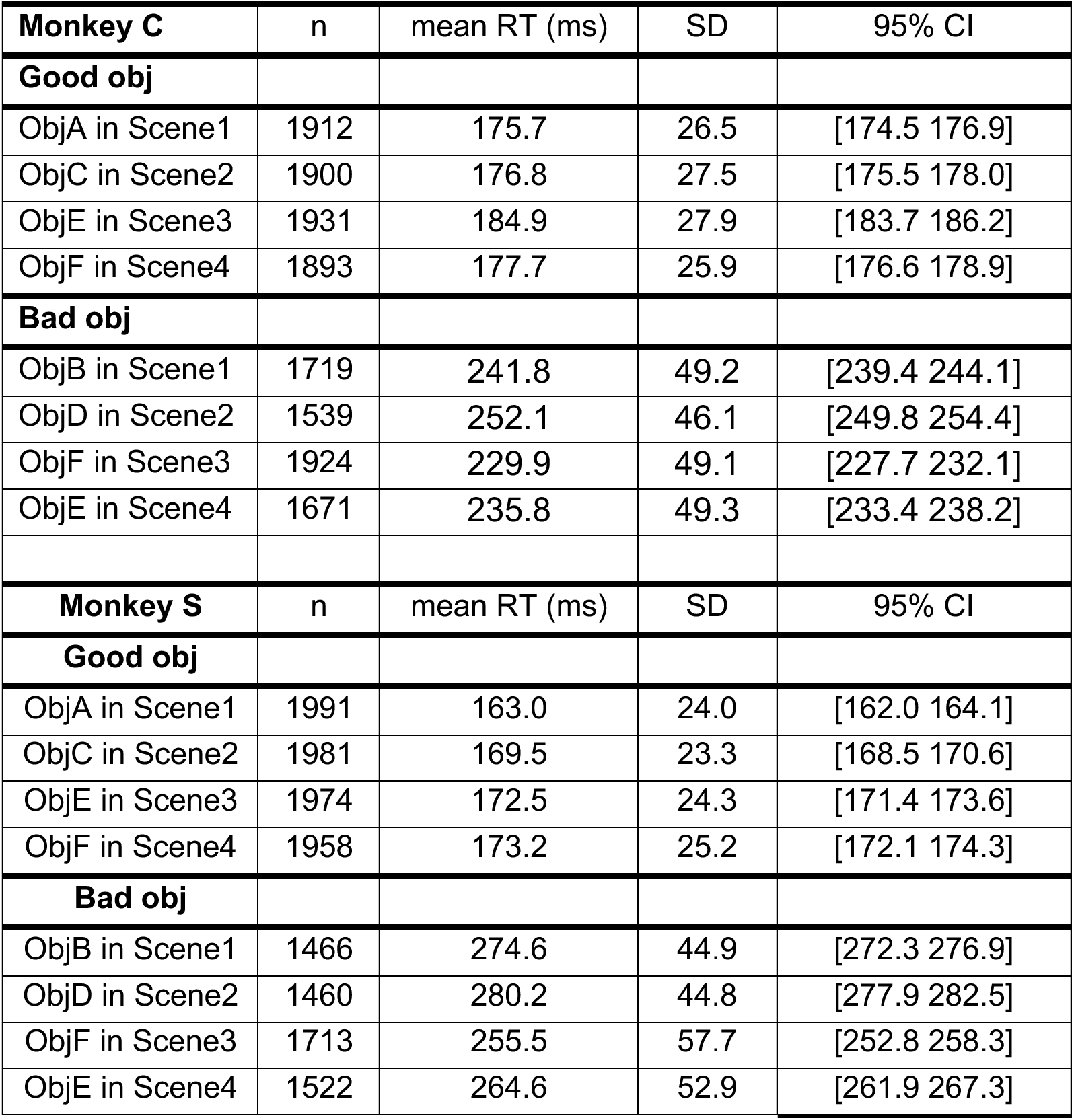
Saccade reaction times in each condition of each monkey.

**Table S2.**
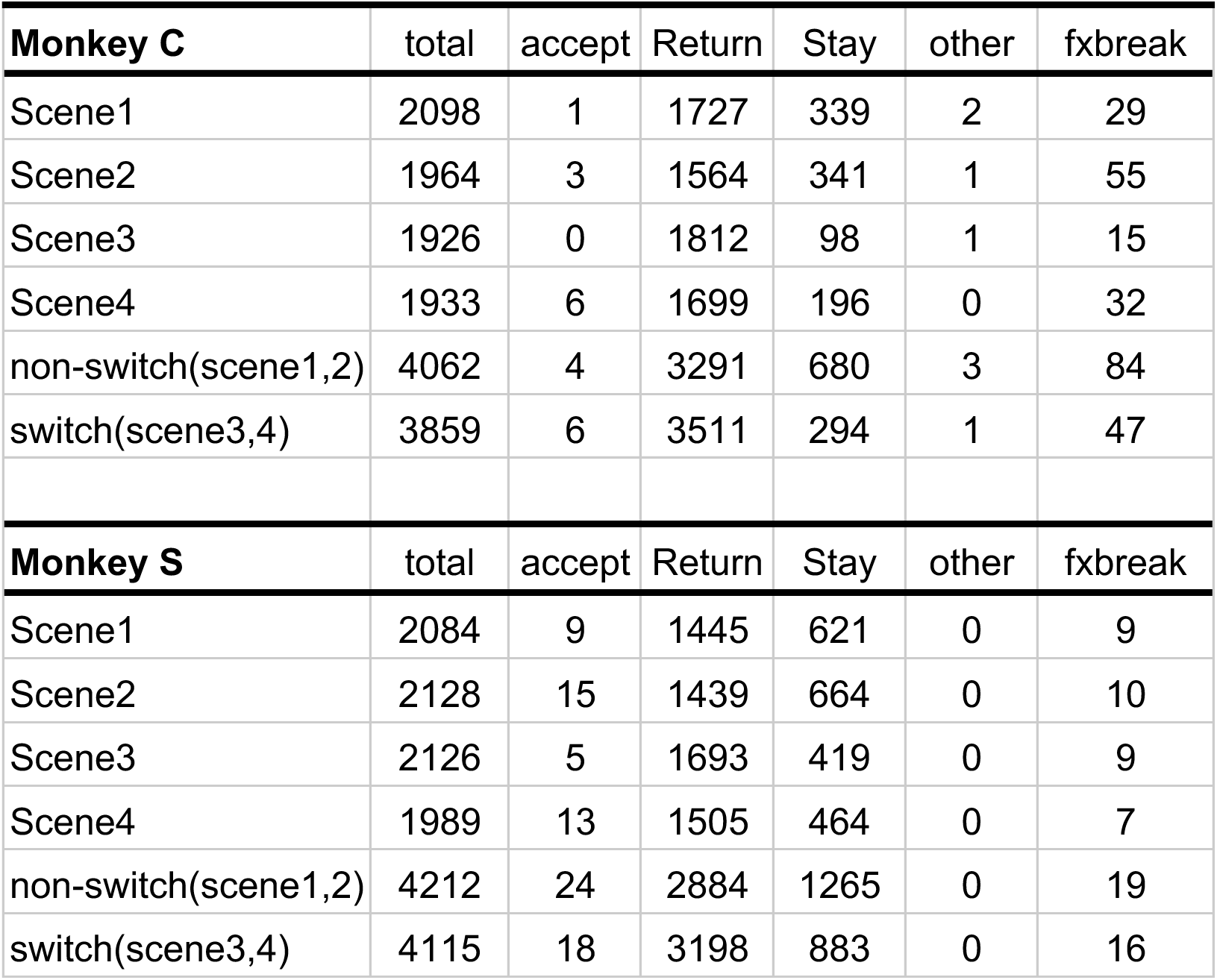
Counts of chosen actions for Bad objects.

**Table S3.**
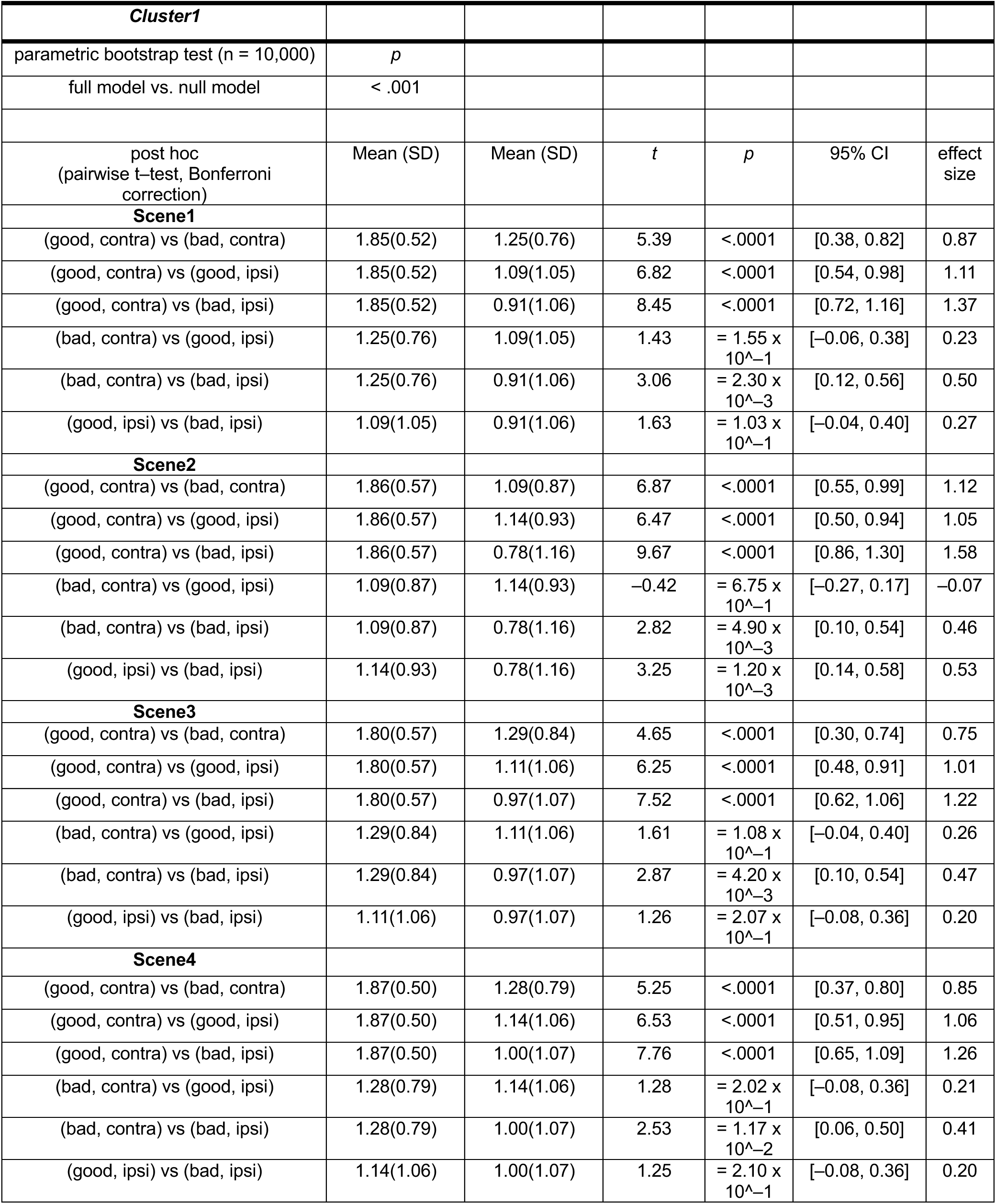
Summary of statistical test to compare the normalized neuronal activity of STN neurons of cluster1 at target onset among conditions during choice task in Figure 3.

**Table S4.**
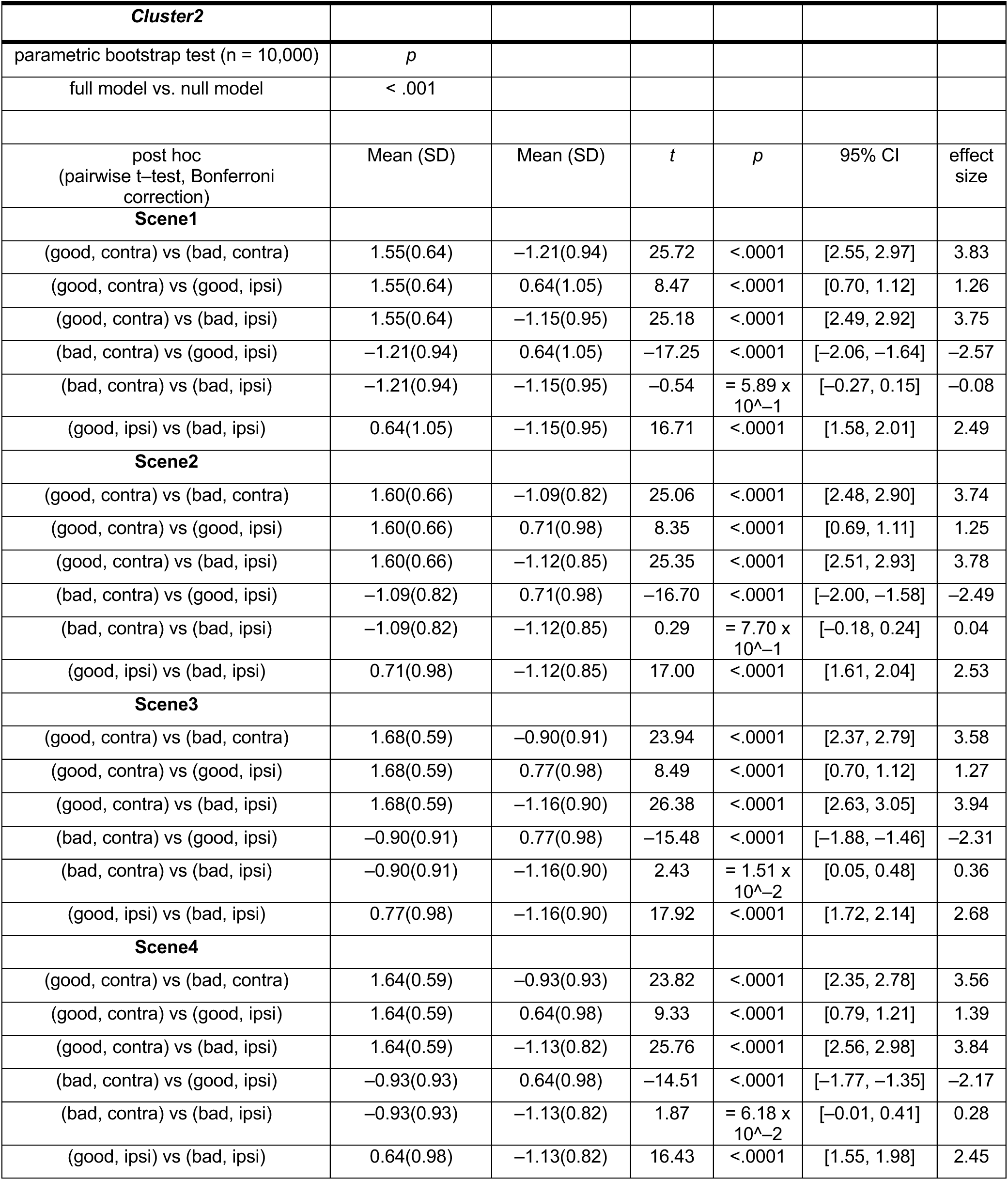
Summary of statistical test to compare the normalized neuronal activity of STN neurons of cluster2 at target onset among conditions during choice task in Figure 3.

**Table S5.**
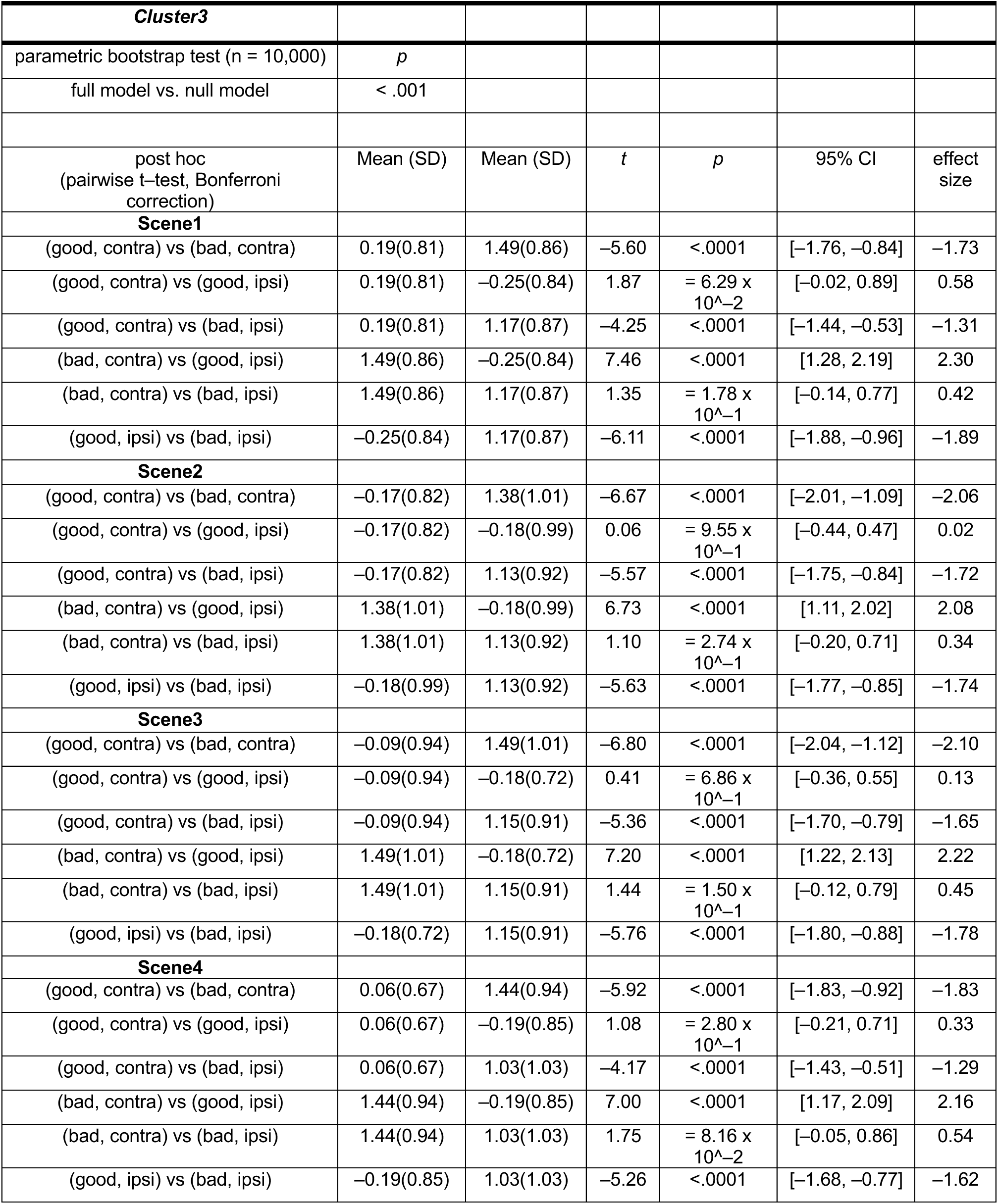
Summary of statistical test to compare the normalized neuronal activity of STN neurons of cluster3 at target onset among conditions during choice task in Figure 3.

**Table S6.**
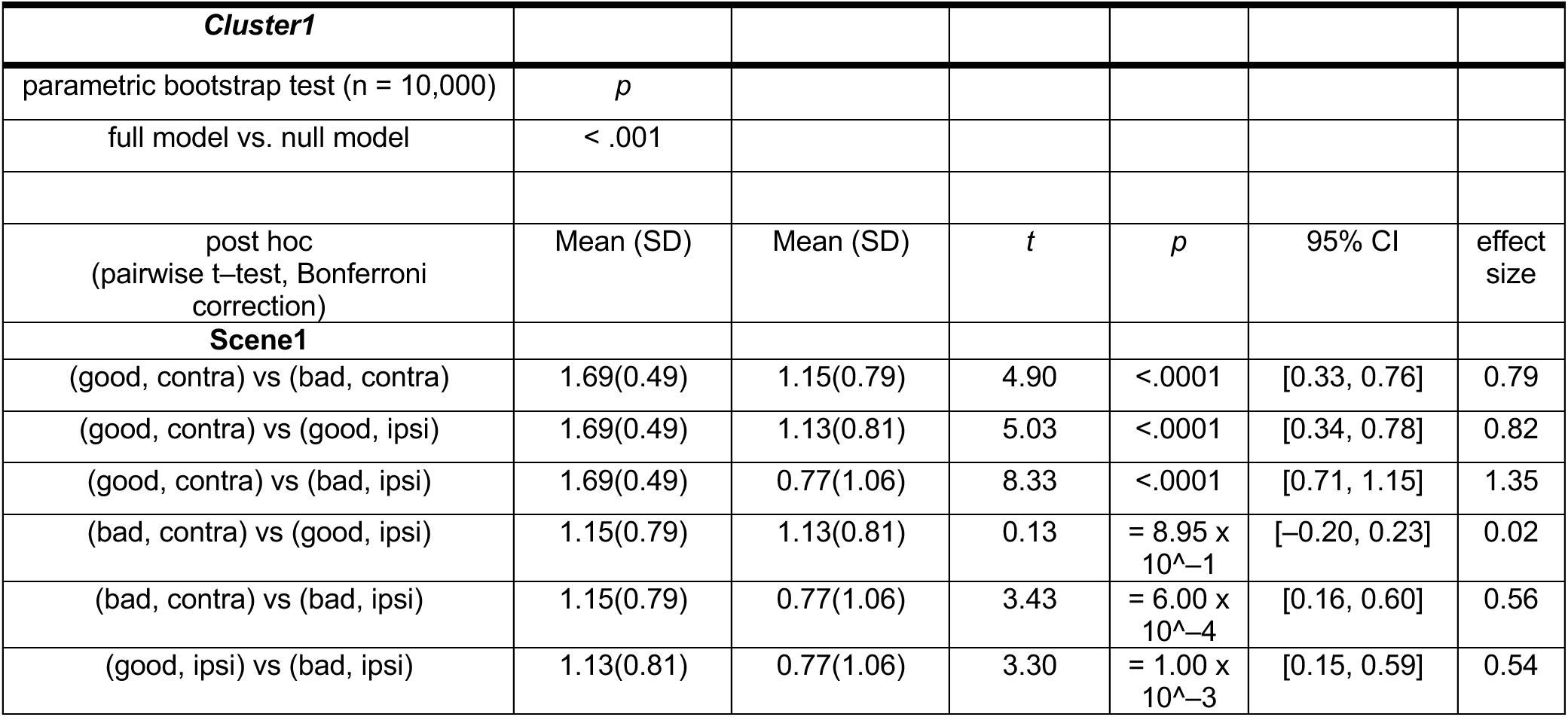
Summary of statistical test to compare the normalized neuronal activity of STN neurons of cluster1 at saccade onset among conditions during choice task in Figure 4.

**Table S7.**
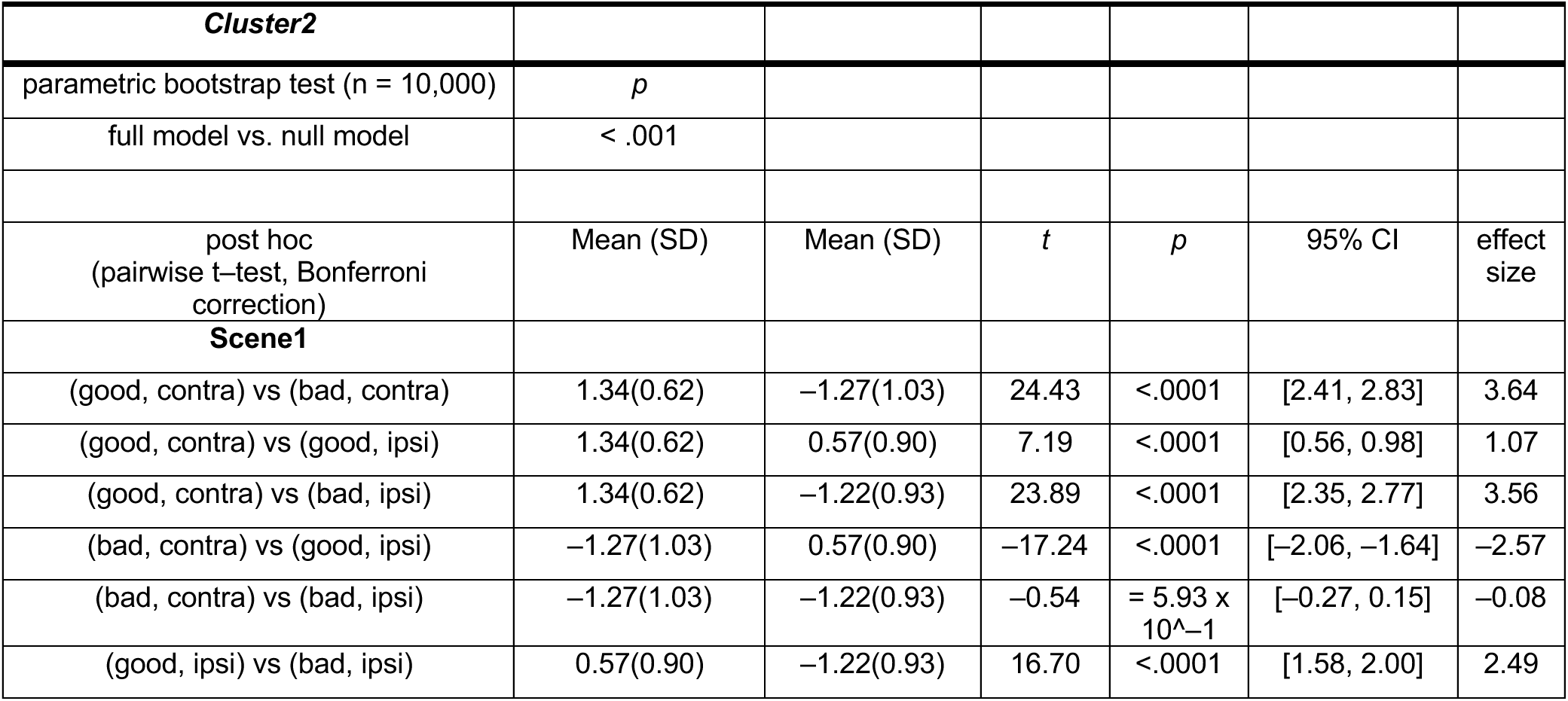
Summary of statistical test to compare the normalized neuronal activity of STN neurons of cluster2 at saccade onset among conditions during choice task in Figure 4.

**Table S8.**
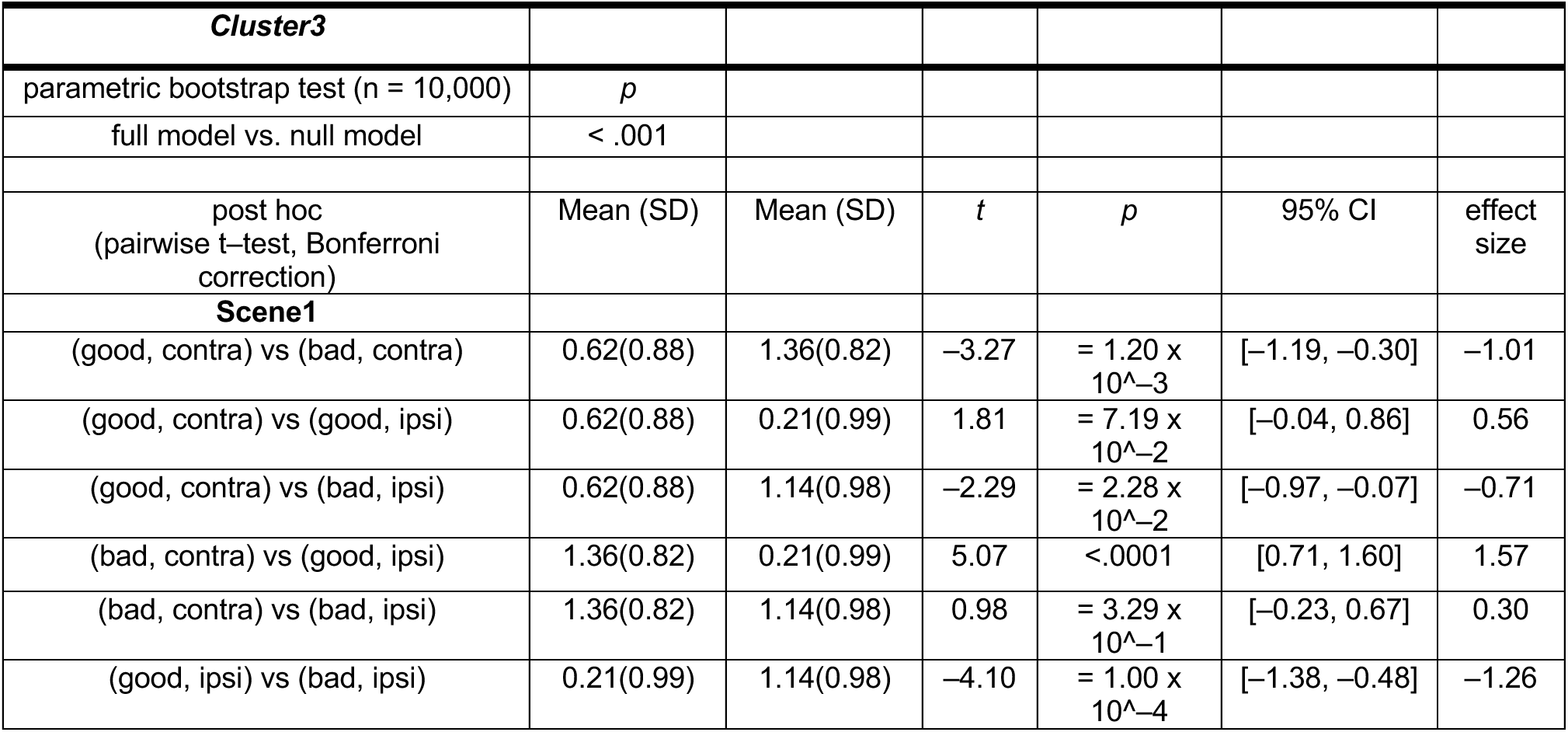
Summary of statistical test to compare the normalized neuronal activity of STN neurons of cluster3 at saccade onset among conditions during choice task in Figure 4.

**Table S9.**
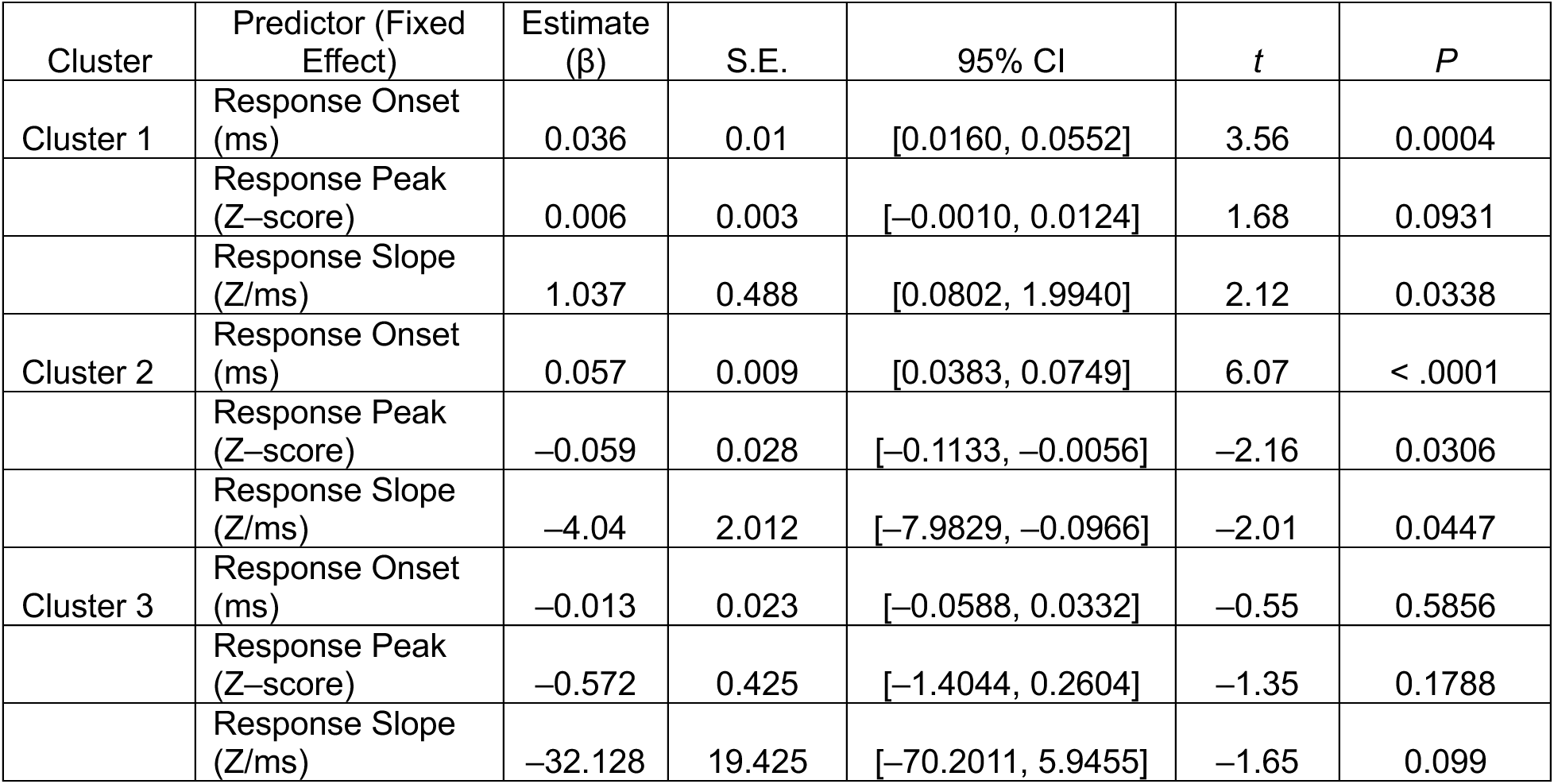
Fixed effects from LMMs models predicting reaction time. Linear mixed–effects models were fitted for each cluster to assess the relationship between neuronal response parameters and reaction time (RT) in Figure S5. Models included NeuronID and MonkeyID as random intercepts.

