## Supplemental materials for "Beyond the Brake: the Subthalamic Nucleus Predominantly Facilitates Action in Non-human Primates"

**Table S1. Saccade reaction times in each condition of each monkey.**

| Monkey C | n | mean RT (ms) | SD | 95% CI |
|---|---|---|---|---|
| **Good obj** | | | | |
| ObjA in Scene1 | 1912 | 175.7 | 26.5 | [174.5 176.9] |
| ObjC in Scene2 | 1900 | 176.8 | 27.5 | [175.5 178.0] |
| ObjE in Scene3 | 1931 | 184.9 | 27.9 | [183.7 186.2] |
| ObjF in Scene4 | 1893 | 177.7 | 25.9 | [176.6 178.9] |
| **Bad obj** | | | | |
| ObjB in Scene1 | 1719 | 241.8 | 49.2 | [239.4 244.1] |
| ObjD in Scene2 | 1539 | 252.1 | 46.1 | [249.8 254.4] |
| ObjF in Scene3 | 1924 | 229.9 | 49.1 | [227.7 232.1] |
| ObjE in Scene4 | 1671 | 235.8 | 49.3 | [233.4 238.2] |
| **Monkey S** | n | mean RT (ms) | SD | 95% CI |
| **Good obj** | | | | |
| ObjA in Scene1 | 1991 | 163.0 | 24.0 | [162.0 164.1] |
| ObjC in Scene2 | 1981 | 169.5 | 23.3 | [168.5 170.6] |
| ObjE in Scene3 | 1974 | 172.5 | 24.3 | [171.4 173.6] |
| ObjF in Scene4 | 1958 | 173.2 | 25.2 | [172.1 174.3] |
| **Bad obj** | | | | |
| ObjB in Scene1 | 1466 | 274.6 | 44.9 | [272.3 276.9] |
| ObjD in Scene2 | 1460 | 280.2 | 44.8 | [277.9 282.5] |
| ObjF in Scene3 | 1713 | 255.5 | 57.7 | [252.8 258.3] |
| ObjE in Scene4 | 1522 | 264.6 | 52.9 | [261.9 267.3] |

**Table S2. Counts of chosen actions for Bad objects**

| **Monkey C** | total | accept | Return | Stay | other | fxbreak |
|---|---|---|---|---|---|---|
| Scene1 | 2098 | 1 | 1727 | 339 | 2 | 29 |
| Scene2 | 1964 | 3 | 1564 | 341 | 1 | 55 |
| Scene3 | 1926 | 0 | 1812 | 98 | 1 | 15 |
| Scene4 | 1933 | 6 | 1699 | 196 | 0 | 32 |
| non-switch(scene1,2) | 4062 | 4 | 3291 | 680 | 3 | 84 |
| switch(scene3,4) | 3859 | 6 | 3511 | 294 | 1 | 47 |
| **Monkey S** | total | accept | Return | Stay | other | fxbreak |
| Scene1 | 2084 | 9 | 1445 | 621 | 0 | 9 |
| Scene2 | 2128 | 15 | 1439 | 664 | 0 | 10 |
| Scene3 | 2126 | 5 | 1693 | 419 | 0 | 9 |
| Scene4 | 1989 | 13 | 1505 | 464 | 0 | 7 |
| non-switch(scene1,2) | 4212 | 24 | 2884 | 1265 | 0 | 19 |
| switch(scene3,4) | 4115 | 18 | 3198 | 883 | 0 | 16 |

**Table S3. Summary of statistical test to compare the normalized neuronal activity of STN neurons of cluster1 at target onset among conditions during choice task in Figure 3.**

| ***Cluster1*** | | | | | | |
|---|---|---|---|---|---|---|
| parametric bootstrap test (n = 10,000) | *p* | | | | | |
| full model vs. null model | < .001 | | | | | |
| post hoc (pairwise t–test, Bonferroni correction) | Mean (SD) | Mean (SD) | *t* | *p* | 95% CI | effect size |
| **Scene1** | | | | | | |
| (good, contra) vs (bad, contra) | 1.85(0.52) | 1.25(0.76) | 5.39 | <.0001 | [0.38, 0.82] | 0.87 |
| (good, contra) vs (good, ipsi) | 1.85(0.52) | 1.09(1.05) | 6.82 | <.0001 | [0.54, 0.98] | 1.11 |
| (good, contra) vs (bad, ipsi) | 1.85(0.52) | 0.91(1.06) | 8.45 | <.0001 | [0.72, 1.16] | 1.37 |
| (bad, contra) vs (good, ipsi) | 1.25(0.76) | 1.09(1.05) | 1.43 | = 1.55 x 10^–1 | [–0.06, 0.38] | 0.23 |
| (bad, contra) vs (bad, ipsi) | 1.25(0.76) | 0.91(1.06) | 3.06 | = 2.30 x 10^–3 | [0.12, 0.56] | 0.50 |
| (good, ipsi) vs (bad, ipsi) | 1.09(1.05) | 0.91(1.06) | 1.63 | = 1.03 x 10^–1 | [–0.04, 0.40] | 0.27 |
| **Scene2** | | | | | | |
| (good, contra) vs (bad, contra) | 1.86(0.57) | 1.09(0.87) | 6.87 | <.0001 | [0.55, 0.99] | 1.12 |
| (good, contra) vs (good, ipsi) | 1.86(0.57) | 1.14(0.93) | 6.47 | <.0001 | [0.50, 0.94] | 1.05 |
| (good, contra) vs (bad, ipsi) | 1.86(0.57) | 0.78(1.16) | 9.67 | <.0001 | [0.86, 1.30] | 1.58 |
| (bad, contra) vs (good, ipsi) | 1.09(0.87) | 1.14(0.93) | –0.42 | = 6.75 x 10^–1 | [–0.27, 0.17] | –0.07 |
| (bad, contra) vs (bad, ipsi) | 1.09(0.87) | 0.78(1.16) | 2.82 | = 4.90 x 10^–3 | [0.10, 0.54] | 0.46 |
| (good, ipsi) vs (bad, ipsi) | 1.14(0.93) | 0.78(1.16) | 3.25 | = 1.20 x 10^–3 | [0.14, 0.58] | 0.53 |
| **Scene3** | | | | | | |
| (good, contra) vs (bad, contra) | 1.80(0.57) | 1.29(0.84) | 4.65 | <.0001 | [0.30, 0.74] | 0.75 |
| (good, contra) vs (good, ipsi) | 1.80(0.57) | 1.11(1.06) | 6.25 | <.0001 | [0.48, 0.91] | 1.01 |
| (good, contra) vs (bad, ipsi) | 1.80(0.57) | 0.97(1.07) | 7.52 | <.0001 | [0.62, 1.06] | 1.22 |
| (bad, contra) vs (good, ipsi) | 1.29(0.84) | 1.11(1.06) | 1.61 | = 1.08 x 10^–1 | [–0.04, 0.40] | 0.26 |
| (bad, contra) vs (bad, ipsi) | 1.29(0.84) | 0.97(1.07) | 2.87 | = 4.20 x 10^–3 | [0.10, 0.54] | 0.47 |
| (good, ipsi) vs (bad, ipsi) | 1.11(1.06) | 0.97(1.07) | 1.26 | = 2.07 x 10^–1 | [–0.08, 0.36] | 0.20 |
| **Scene4** | | | | | | |
| (good, contra) vs (bad, contra) | 1.87(0.50) | 1.28(0.79) | 5.25 | <.0001 | [0.37, 0.80] | 0.85 |
| (good, contra) vs (good, ipsi) | 1.87(0.50) | 1.14(1.06) | 6.53 | <.0001 | [0.51, 0.95] | 1.06 |
| (good, contra) vs (bad, ipsi) | 1.87(0.50) | 1.00(1.07) | 7.76 | <.0001 | [0.65, 1.09] | 1.26 |
| (bad, contra) vs (good, ipsi) | 1.28(0.79) | 1.14(1.06) | 1.28 | = 2.02 x 10^–1 | [–0.08, 0.36] | 0.21 |
| (bad, contra) vs (bad, ipsi) | 1.28(0.79) | 1.00(1.07) | 2.53 | = 1.17 x 10^–2 | [0.06, 0.50] | 0.41 |
| (good, ipsi) vs (bad, ipsi) | 1.14(1.06) | 1.00(1.07) | 1.25 | = 2.10 x 10^–1 | [–0.08, 0.36] | 0.20 |

**Table S4. Summary of statistical test to compare the normalized neuronal activity of STN neurons of cluster2 at target onset among conditions during choice task in Figure 3.**

| ***Cluster2*** | | | | | | |
|---|---|---|---|---|---|---|
| parametric bootstrap test (n = 10,000) | *p* | | | | | |
| full model vs. null model | < .001 | | | | | |
| post hoc (pairwise t–test, Bonferroni correction) | Mean (SD) | Mean (SD) | *t* | *p* | 95% CI | effect size |
| **Scene1** | | | | | | |
| (good, contra) vs (bad, contra) | 1.55(0.64) | –1.21(0.94) | 25.72 | <.0001 | [2.55, 2.97] | 3.83 |
| (good, contra) vs (good, ipsi) | 1.55(0.64) | 0.64(1.05) | 8.47 | <.0001 | [0.70, 1.12] | 1.26 |
| (good, contra) vs (bad, ipsi) | 1.55(0.64) | –1.15(0.95) | 25.18 | <.0001 | [2.49, 2.92] | 3.75 |
| (bad, contra) vs (good, ipsi) | –1.21(0.94) | 0.64(1.05) | –17.25 | <.0001 | [–2.06, –1.64] | –2.57 |
| (bad, contra) vs (bad, ipsi) | –1.21(0.94) | –1.15(0.95) | –0.54 | = 5.89 x 10^–1 | [–0.27, 0.15] | –0.08 |
| (good, ipsi) vs (bad, ipsi) | 0.64(1.05) | –1.15(0.95) | 16.71 | <.0001 | [1.58, 2.01] | 2.49 |
| **Scene2** | | | | | | |
| (good, contra) vs (bad, contra) | 1.60(0.66) | –1.09(0.82) | 25.06 | <.0001 | [2.48, 2.90] | 3.74 |
| (good, contra) vs (good, ipsi) | 1.60(0.66) | 0.71(0.98) | 8.35 | <.0001 | [0.69, 1.11] | 1.25 |
| (good, contra) vs (bad, ipsi) | 1.60(0.66) | –1.12(0.85) | 25.35 | <.0001 | [2.51, 2.93] | 3.78 |
| (bad, contra) vs (good, ipsi) | –1.09(0.82) | 0.71(0.98) | –16.70 | <.0001 | [–2.00, –1.58] | –2.49 |
| (bad, contra) vs (bad, ipsi) | –1.09(0.82) | –1.12(0.85) | 0.29 | = 7.70 x 10^–1 | [–0.18, 0.24] | 0.04 |
| (good, ipsi) vs (bad, ipsi) | 0.71(0.98) | –1.12(0.85) | 17.00 | <.0001 | [1.61, 2.04] | 2.53 |
| **Scene3** | | | | | | |
| (good, contra) vs (bad, contra) | 1.68(0.59) | –0.90(0.91) | 23.94 | <.0001 | [2.37, 2.79] | 3.58 |
| (good, contra) vs (good, ipsi) | 1.68(0.59) | 0.77(0.98) | 8.49 | <.0001 | [0.70, 1.12] | 1.27 |
| (good, contra) vs (bad, ipsi) | 1.68(0.59) | –1.16(0.90) | 26.38 | <.0001 | [2.63, 3.05] | 3.94 |
| (bad, contra) vs (good, ipsi) | –0.90(0.91) | 0.77(0.98) | –15.48 | <.0001 | [–1.88, –1.46] | –2.31 |
| (bad, contra) vs (bad, ipsi) | –0.90(0.91) | –1.16(0.90) | 2.43 | = 1.51 x 10^–2 | [0.05, 0.48] | 0.36 |
| (good, ipsi) vs (bad, ipsi) | 0.77(0.98) | –1.16(0.90) | 17.92 | <.0001 | [1.72, 2.14] | 2.68 |
| **Scene4** | | | | | | |
| (good, contra) vs (bad, contra) | 1.64(0.59) | –0.93(0.93) | 23.82 | <.0001 | [2.35, 2.78] | 3.56 |
| (good, contra) vs (good, ipsi) | 1.64(0.59) | 0.64(0.98) | 9.33 | <.0001 | [0.79, 1.21] | 1.39 |
| (good, contra) vs (bad, ipsi) | 1.64(0.59) | –1.13(0.82) | 25.76 | <.0001 | [2.56, 2.98] | 3.84 |
| (bad, contra) vs (good, ipsi) | –0.93(0.93) | 0.64(0.98) | –14.51 | <.0001 | [–1.77, –1.35] | –2.17 |
| (bad, contra) vs (bad, ipsi) | –0.93(0.93) | –1.13(0.82) | 1.87 | = 6.18 x 10^–2 | [–0.01, 0.41] | 0.28 |
| (good, ipsi) vs (bad, ipsi) | 0.64(0.98) | –1.13(0.82) | 16.43 | <.0001 | [1.55, 1.98] | 2.45 |

**Table S5. Summary of statistical test to compare the normalized neuronal activity of STN neurons of cluster3 at target onset among conditions during choice task in Figure 3.**

| ***Cluster3*** | | | | | | |
|---|---|---|---|---|---|---|
| parametric bootstrap test (n = 10,000) | *p* | | | | | |
| full model vs. null model | < .001 | | | | | |
| post hoc (pairwise t–test, Bonferroni correction) | Mean (SD) | Mean (SD) | *t* | *p* | 95% CI | effect size |
| **Scene1** | | | | | | |
| (good, contra) vs (bad, contra) | 0.19(0.81) | 1.49(0.86) | –5.60 | <.0001 | [–1.76, –0.84] | –1.73 |
| (good, contra) vs (good, ipsi) | 0.19(0.81) | –0.25(0.84) | 1.87 | = 6.29 x 10^–2 | [–0.02, 0.89] | 0.58 |
| (good, contra) vs (bad, ipsi) | 0.19(0.81) | 1.17(0.87) | –4.25 | <.0001 | [–1.44, –0.53] | –1.31 |
| (bad, contra) vs (good, ipsi) | 1.49(0.86) | –0.25(0.84) | 7.46 | <.0001 | [1.28, 2.19] | 2.30 |
| (bad, contra) vs (bad, ipsi) | 1.49(0.86) | 1.17(0.87) | 1.35 | = 1.78 x 10^–1 | [–0.14, 0.77] | 0.42 |
| (good, ipsi) vs (bad, ipsi) | –0.25(0.84) | 1.17(0.87) | –6.11 | <.0001 | [–1.88, –0.96] | –1.89 |
| **Scene2** | | | | | | |
| (good, contra) vs (bad, contra) | –0.17(0.82) | 1.38(1.01) | –6.67 | <.0001 | [–2.01, –1.09] | –2.06 |
| (good, contra) vs (good, ipsi) | –0.17(0.82) | –0.18(0.99) | 0.06 | = 9.55 x 10^–1 | [–0.44, 0.47] | 0.02 |
| (good, contra) vs (bad, ipsi) | –0.17(0.82) | 1.13(0.92) | –5.57 | <.0001 | [–1.75, –0.84] | –1.72 |
| (bad, contra) vs (good, ipsi) | 1.38(1.01) | –0.18(0.99) | 6.73 | <.0001 | [1.11, 2.02] | 2.08 |
| (bad, contra) vs (bad, ipsi) | 1.38(1.01) | 1.13(0.92) | 1.10 | = 2.74 x 10^–1 | [–0.20, 0.71] | 0.34 |
| (good, ipsi) vs (bad, ipsi) | –0.18(0.99) | 1.13(0.92) | –5.63 | <.0001 | [–1.77, –0.85] | –1.74 |
| **Scene3** | | | | | | |
| (good, contra) vs (bad, contra) | –0.09(0.94) | 1.49(1.01) | –6.80 | <.0001 | [–2.04, –1.12] | –2.10 |
| (good, contra) vs (good, ipsi) | –0.09(0.94) | –0.18(0.72) | 0.41 | = 6.86 x 10^–1 | [–0.36, 0.55] | 0.13 |
| (good, contra) vs (bad, ipsi) | –0.09(0.94) | 1.15(0.91) | –5.36 | <.0001 | [–1.70, –0.79] | –1.65 |
| (bad, contra) vs (good, ipsi) | 1.49(1.01) | –0.18(0.72) | 7.20 | <.0001 | [1.22, 2.13] | 2.22 |
| (bad, contra) vs (bad, ipsi) | 1.49(1.01) | 1.15(0.91) | 1.44 | = 1.50 x 10^–1 | [–0.12, 0.79] | 0.45 |
| (good, ipsi) vs (bad, ipsi) | –0.18(0.72) | 1.15(0.91) | –5.76 | <.0001 | [–1.80, –0.88] | –1.78 |
| **Scene4** | | | | | | |
| (good, contra) vs (bad, contra) | 0.06(0.67) | 1.44(0.94) | –5.92 | <.0001 | [–1.83, –0.92] | –1.83 |
| (good, contra) vs (good, ipsi) | 0.06(0.67) | –0.19(0.85) | 1.08 | = 2.80 x 10^–1 | [–0.21, 0.71] | 0.33 |
| (good, contra) vs (bad, ipsi) | 0.06(0.67) | 1.03(1.03) | –4.17 | <.0001 | [–1.43, –0.51] | –1.29 |
| (bad, contra) vs (good, ipsi) | 1.44(0.94) | –0.19(0.85) | 7.00 | <.0001 | [1.17, 2.09] | 2.16 |
| (bad, contra) vs (bad, ipsi) | 1.44(0.94) | 1.03(1.03) | 1.75 | = 8.16 x 10^–2 | [–0.05, 0.86] | 0.54 |
| (good, ipsi) vs (bad, ipsi) | –0.19(0.85) | 1.03(1.03) | –5.26 | <.0001 | [–1.68, –0.77] | –1.62 |

**Table S6. Summary of statistical test to compare the normalized neuronal activity of STN neurons of cluster1 at saccade onset among conditions during choice task in Figure 4.**

| ***Cluster1*** | | | | | | |
|---|---|---|---|---|---|---|
| parametric bootstrap test (n = 10,000) | *p* | | | | | |
| full model vs. null model | < .001 | | | | | |
| post hoc (pairwise t–test, Bonferroni correction) | Mean (SD) | Mean (SD) | *t* | *p* | 95% CI | effect size |
| **Scene1** | | | | | | |
| (good, contra) vs (bad, contra) | 1.69(0.49) | 1.15(0.79) | 4.90 | <.0001 | [0.33, 0.76] | 0.79 |
| (good, contra) vs (good, ipsi) | 1.69(0.49) | 1.13(0.81) | 5.03 | <.0001 | [0.34, 0.78] | 0.82 |
| (good, contra) vs (bad, ipsi) | 1.69(0.49) | 0.77(1.06) | 8.33 | <.0001 | [0.71, 1.15] | 1.35 |
| (bad, contra) vs (good, ipsi) | 1.15(0.79) | 1.13(0.81) | 0.13 | = 8.95 x 10^–1 | [–0.20, 0.23] | 0.02 |
| (bad, contra) vs (bad, ipsi) | 1.15(0.79) | 0.77(1.06) | 3.43 | = 6.00 x 10^–4 | [0.16, 0.60] | 0.56 |
| (good, ipsi) vs (bad, ipsi) | 1.13(0.81) | 0.77(1.06) | 3.30 | = 1.00 x 10^–3 | [0.15, 0.59] | 0.54 |

**Table S7. Summary of statistical test to compare the normalized neuronal activity of STN neurons of cluster2 at saccade onset among conditions during choice task in Figure 4.**

| ***Cluster2*** | | | | | | |
|---|---|---|---|---|---|---|
| parametric bootstrap test (n = 10,000) | *p* | | | | | |
| full model vs. null model | < .001 | | | | | |
| post hoc (pairwise t–test, Bonferroni correction) | Mean (SD) | Mean (SD) | *t* | *p* | 95% CI | effect size |
| **Scene1** | | | | | | |
| (good, contra) vs (bad, contra) | 1.34(0.62) | –1.27(1.03) | 24.43 | <.0001 | [2.41, 2.83] | 3.64 |
| (good, contra) vs (good, ipsi) | 1.34(0.62) | 0.57(0.90) | 7.19 | <.0001 | [0.56, 0.98] | 1.07 |
| (good, contra) vs (bad, ipsi) | 1.34(0.62) | –1.22(0.93) | 23.89 | <.0001 | [2.35, 2.77] | 3.56 |
| (bad, contra) vs (good, ipsi) | –1.27(1.03) | 0.57(0.90) | –17.24 | <.0001 | [–2.06, –1.64] | –2.57 |
| (bad, contra) vs (bad, ipsi) | –1.27(1.03) | –1.22(0.93) | –0.54 | = 5.93 x 10^–1 | [–0.27, 0.15] | –0.08 |
| (good, ipsi) vs (bad, ipsi) | 0.57(0.90) | –1.22(0.93) | 16.70 | <.0001 | [1.58, 2.00] | 2.49 |

**Table S8. Summary of statistical test to compare the normalized neuronal activity of STN neurons of cluster3 at saccade onset among conditions during choice task in Figure 4.**

| ***Cluster3*** | | | | | | |
|---|---|---|---|---|---|---|
| parametric bootstrap test (n = 10,000) | *p* | | | | | |
| full model vs. null model | < .001 | | | | | |
| post hoc (pairwise t–test, Bonferroni correction) | Mean (SD) | Mean (SD) | *t* | *p* | 95% CI | effect size |
| **Scene1** | | | | | | |
| (good, contra) vs (bad, contra) | 0.62(0.88) | 1.36(0.82) | –3.27 | = 1.20 x 10^–3 | [–1.19, –0.30] | –1.01 |
| (good, contra) vs (good, ipsi) | 0.62(0.88) | 0.21(0.99) | 1.81 | = 7.19 x 10^–2 | [–0.04, 0.86] | 0.56 |
| (good, contra) vs (bad, ipsi) | 0.62(0.88) | 1.14(0.98) | –2.29 | = 2.28 x 10^–2 | [–0.97, –0.07] | –0.71 |
| (bad, contra) vs (good, ipsi) | 1.36(0.82) | 0.21(0.99) | 5.07 | <.0001 | [0.71, 1.60] | 1.57 |
| (bad, contra) vs (bad, ipsi) | 1.36(0.82) | 1.14(0.98) | 0.98 | = 3.29 x 10^–1 | [–0.23, 0.67] | 0.30 |
| (good, ipsi) vs (bad, ipsi) | 0.21(0.99) | 1.14(0.98) | –4.10 | = 1.00 x 10^–4 | [–1.38, –0.48] | –1.26 |

**Table S9. Fixed effects from LMMs models predicting reaction time. Linear mixed–effects models were fitted for each cluster to assess the relationship between neuronal response parameters and reaction time (RT). Models included NeuronID and MonkeyID as random intercepts.**

| Cluster | Predictor (Fixed Effect) | Estimate (β) | S.E. | 95% CI | *t* | *P* |
|---|---|---|---|---|---|---|
| Cluster 1 | Response Onset (ms) | 0.036 | 0.01 | [0.0160, 0.0552] | 3.56 | 0.0004 |
| | Response Peak (Z–score) | 0.006 | 0.003 | [–0.0010, 0.0124] | 1.68 | 0.0931 |
| | Response Slope (Z/ms) | 1.037 | 0.488 | [0.0802, 1.9940] | 2.12 | 0.0338 |
| Cluster 2 | Response Onset (ms) | 0.057 | 0.009 | [0.0383, 0.0749] | 6.07 | < .0001 |
| | Response Peak (Z–score) | –0.059 | 0.028 | [–0.1133, –0.0056] | –2.16 | 0.0306 |
| | Response Slope (Z/ms) | –4.04 | 2.012 | [–7.9829, –0.0966] | –2.01 | 0.0447 |
| Cluster 3 | Response Onset (ms) | –0.013 | 0.023 | [–0.0588, 0.0332] | –0.55 | 0.5856 |
| | Response Peak (Z–score) | –0.572 | 0.425 | [–1.4044, 0.2604] | –1.35 | 0.1788 |
| | Response Slope (Z/ms) | –32.128 | 19.425 | [–70.2011, 5.9455] | –1.65 | 0.099 |
